# Dynamic regulation of JAK-STAT signaling through the prolactin receptor predicted by computational modeling

**DOI:** 10.1101/2020.02.14.949321

**Authors:** Ryland D. Mortlock, Senta K. Georgia, Stacey D. Finley

## Abstract

**Introduction:** Hormones signal through various receptors and cascades of biochemical reactions to expand beta cell mass during pregnancy. Harnessing this phenomenon to treat beta cell dysfunction requires quantitative understanding of the signaling at the molecular level. This study explores how different regulatory elements impact JAK-STAT signaling through the prolactin receptor in pancreatic beta cells.

**Methods:** A mechanistic computational model was constructed to describe the key reactions and molecular species involved in JAK-STAT signaling in response to the hormone prolactin. The effect of including and excluding different regulatory modules in the model structure was explored through ensemble modeling. A Bayesian approach for likelihood estimation was used to parametrize the model to experimental data from the literature.

**Results:** Receptor upregulation, combined with either inhibition by SOCS proteins, receptor internalization, or both, was required to obtain STAT5 dynamics matching experimental results for INS-1 cells treated with prolactin. Multiple model structures could fit the experimental data, and key findings were conserved across model structures, including faster dimerization and nuclear import rates of STAT5B compared to STAT5A. The model was validated using experimental data from rat primary beta cells not used in parameter estimation. Probing the fitted, validated model revealed possible strategies to modulate STAT5 signaling.

**Conclusions:** JAK-STAT signaling must be tightly controlled to obtain the biphasic response in STAT5 activation seen experimentally. Receptor up-regulation, combined with SOCS inhibition, receptor internalization, or both is required to match experimental data. Modulating reactions upstream in the signaling can enhance STAT5 activation to increase beta cell mass.

## 1 INTRODUCTION

Metabolic diseases impair the body’s ability to properly convert nutrients into energy. Diabetes is a particularly harmful metabolic disease that affects over 30 million people in the United States alone.^51^ In cases of Type 1 diabetes, the body is unable to produce insulin, a key hormone that regulates the transport of glucose from the blood to the cells where it is used to produce energy. Patients with Type 2 or gestational diabetes can produce some insulin, but not enough to properly regulate blood glucose levels. Recent advances in the study of pancreatic beta cells, the cells that produce and secrete insulin, have shed light on the body’s ability to adapt in response to changes in metabolic demand.^36^ For example, in cases of high insulin demand, such as pregnancy or obesity, the body responds by increasing beta cell mass in the pancreas. In fact, studies have shown that over the approximately 20-day time course of pregnancy in mice, pancreatic beta cells both replicate and grow in size, resulting in an increased beta cell mass.^36^ This phenomenon could potentially be harnessed to increase the number of functioning beta cells in diabetes patients.

Beta cell compensation precedes insulin compensation and is driven by signaling through the prolactin receptor^4,17,23,47^ (PRLR), which is closely related to the growth hormone receptor. Signaling by placental lactogens through PRLR engages the JAK-STAT signaling cascade.^34^ Specifically, Janus Kinase 2 (JAK2) is constitutively associated with the PRLR^7,15,38^ and once the JAK2 kinase is activated, it recruits and phosphorylates Signal Transducer and Activator of Transcription 5 (STAT5). STAT5 regulates the expression of several target genes in the nucleus, including genes related to the cell cycle^18,41^ and apoptosis.^19,24,46^ Although initial discoveries were made in rodent models, human prolactin has been shown to protect human beta cells from apoptosis as well.^46^

In this work, we investigate the mechanisms by which the pregnancy-related hormone prolactin (PRL) drives JAK-STAT signaling in pancreatic beta cells using a mathematical model of the signaling pathway. Mathematical models have been used to elucidate the balance between replication and apoptosis in beta cells^29^, but no molecular-detailed computational model exists for the beta cell response to pregnancy. Here, we use a systems biology approach to quantitatively analyze the beta cell response to hormone stimulation. We have constructed a computational model, calibrated the model with experimental data, and validated it by predicting new data not used for parameter estimation. During model construction, we explored the effect of different model structures on the predicted signaling dynamics through an approach known as ensemble modeling.^25,31^ After validating the model, we analyzed how changes in kinetic parameters and initial values can lead to greater STAT5 activation.

In particular, mathematical modeling is used to explore the effects of various regulatory mechanisms that control signaling. Experimental data shows that when insulin-secreting cells of the INS-1 cell line are treated with a constant concentration of PRL *in vitro*, the amount of phosphorylated STAT5 has multiple peaks within six hours of stimulation.^10,11^ The presence of these peaks is influenced by Suppressors of Cytokine Signaling (SOCS) genes, which are transcribed in response to STAT signaling and exert negative feedback on the system. Modeling the cytokine IFN-γ in liver cells, Yamada *et al*. found that the presence of a nuclear phosphatase, in addition to SOCS negative feedback, are sufficient to cause a decrease in phosphorylated STAT after the initial peak, leading to multiple peaks in phosphorylated STAT dimer in the nucleus.^48^ In addition, JAK-STAT signaling through the prolactin receptor (PRLR) has been shown to include positive feedback as nuclear STAT5 promotes transcription of PRLR mRNA.^20,27,33,35^ We hypothesize that positive feedback could play a role in explaining the initial peak, subsequent decline, then prolonged activation of STAT5 activity in INS-1 cells discovered by Brelje and colleagues. Although these regulatory mechanisms significantly influence beta cell signaling, no model to our knowledge explores the interplay between SOCS feedback and positive regulation of PRL signaling. Therefore, we built upon the work of Yamada and colleagues to create a computational model of JAK-STAT signaling in pancreatic beta cells through PRLR. We applied the model to investigate the influence of these regulation schemes, individually and in combination, and found that model structures that include both positive and negative regulation produce multiple peaks in STAT5 phosphorylation within a tight range of parameter values. By fitting to experimental data using a Bayesian approach for likelihood estimation of parameter values, we show that the model can simultaneously determine the rates of STAT5 phosphorylation and nuclear translocation. The model predicts a faster dimerization and nuclear import rate for STAT5B dimers than STAT5A which can explain their different activation profiles observed experimentally. Our experimentally-derived mathematical model provides a quantitative framework needed to better understand signaling that mediates beta cell increase.

## 2 METHODS

### Model construction

A mathematical model was constructed to describe the reaction kinetics of the Janus kinase 2 (JAK2) and signal transducer and activator of transcription 5 (STAT5) signaling in pancreatic beta cells. The model is comprised of ordinary differential equations, which describe how the concentrations of the molecular species in the reaction network evolve over time. Our model builds on the reactions and kinetic parameters from the work of Yamada, *et al*., who modeled control mechanisms of the JAK-STAT pathway in response to interferon-γ (IFN- γ) signaling.^48^ The model was adapted to include 2:1 ligand to receptor stoichiometry, which has been shown for the binding of prolactin (PRL) to the prolactin receptor (PRLR).^7,15^ The receptor is assumed to be pre-associated with JAK2 (represented by the species RJ) since JAK2 is constitutively associated with the prolactin receptor.^7,15,38^ Once two RJ complexes are bound to one PRL hormone, the complex becomes activated. The receptor complex RJ has degradation and synthesis rates corresponding to a half-life on the cell membrane of 45 minutes.^10^ Once the ligand is bound, the receptor has a higher degradation rate, which represents internalization of the ligated receptor to the endosome.^1,7,10^

The activated receptor complex binds to the cytosolic form of STAT5 reversibly, and once bound, releases a phosphorylated form of STAT5 due to the kinase activity of JAK2. The pSTAT5 molecules dimerize in the cytosol and are transported into the nucleus. Three phosphatases are included in the model, which serve to attenuate the signaling after initial ligand binding: SH2 domain-containing tyrosine phosphatase 2 (SHP-2) dephosphorylates the activated receptor-JAK complex, and phosphatases in the cytosol and nucleus (termed PPX and PPN, respectively) dephosphorylate STAT5 species^48^. The phosphatase action is a form of negative feedback shown to be necessary for attenuation of STAT activation.^48^ pSTAT molecules are shuttled out of the nucleus when they are not dimerized with another molecule. The phosphorylated STAT5 dimer promotes transcription of several target genes once in the nucleus. Specifically, we include suppressors of cytokine signaling (SOCS), the prolactin receptor, and the antiapoptotic protein Bcl-xL as STAT5 targets. It has been shown that suppressors of cytokine signaling (SOCS) proteins bind competitively to the receptor JAK complexes and also target the receptors for ubiq-uitination-based degradation.^7,49^ These mechanisms were incorporated in the model rather than the non-competitive binding used by Yamada, *et al*.^48^

STAT5 dimers promote transcription of mRNA for the prolactin receptor. This has been shown *in vitro* in INS-1 cells^20^ and *in vivo* during pregnancy in mice. This positive feedback mechanism may play a role in the islet response to pregnancy^20^ and has not been explored computationally before. The phosphorylated STAT5 dimer in the nucleus also promotes transcription of cell-cycle genes such as cyclin D proteins^18,41^ and anti-apoptotic species such as Bcl-family proteins^19,46^. We included a module for the STAT5-mediated transcription and translation of the response protein Bcl-xL. A full list of reactions is included in the supplementary File S1. MATLAB was used to carry out model simulations, and statistical analyses of the simulated results were performed using R statistical computing language.^45^ All code including MATLAB files, SBML files, and R scripts are publicly available at: https://github.com/ryland-mortlock/Modeling-JAK-STAT-Regulation-Through-Prolactin-Receptor.

### Ensemble modeling

The three optional modules (Fig. 1) were included or excluded from the core model. The induction of SOCS in response to STAT5 activation and its subsequent negative feedback on JAK-STAT signaling was the first optional module. The positive regulation due to up-regulation of the PRL receptor in response to activated STAT5 was the second optional module. The third optional module was receptor internalization, as represented by an enhanced degradation rate for lig-and-bound receptors. The three optional modules were included in different combinations to produce eight possible model structures.

**Figure 1:**
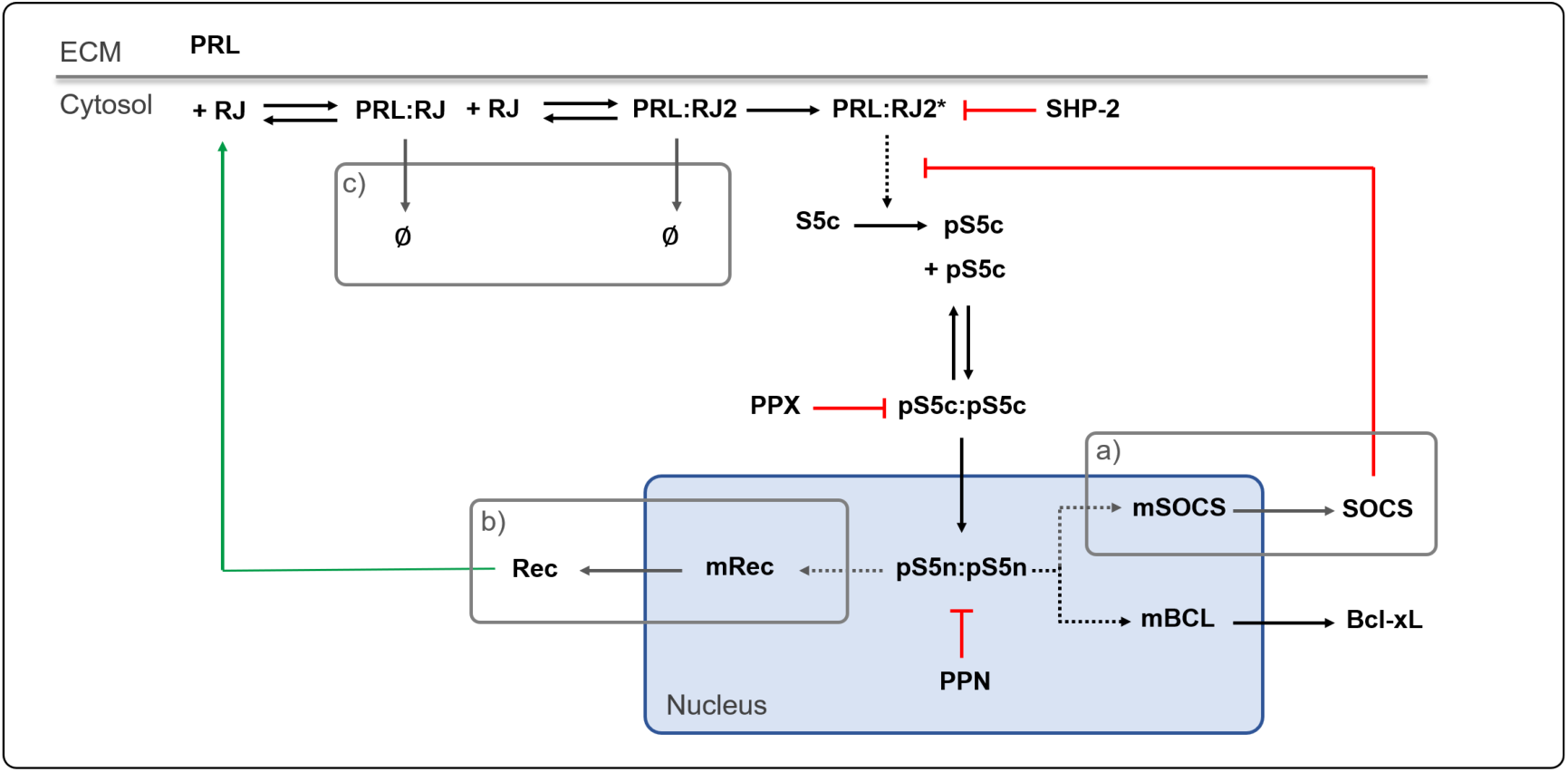
Model Schematic of JAK-STAT signaling in pancreatic beta cells. PRL binds to the PRLr:JAK2 complex (RJ), which induces receptor dimerization and activation by JAK2 kinase activity. The activated receptor PRL:RJ2* phosphorylates STAT5, which dimerizes and transports into the nucleus, where it promotes transcription of target genes. Phosphatases attenuate the signaling at the membrane (SHP-2), in the cytosol (PPX), and in the nucleus (PPN). Signaling modules for ensemble modeling include **a)** STAT5-induced SOCS negative feedback, **b)** STAT5-induced receptor up-regulation, and **c)** ligand-induced receptor internalization. Green indicates positive feedback; red indicates inhibition of signaling. *ECM*, extracellular matrix.

For each model structure, 100,000 Monte Carlo simulations were performed by sampling all free parameters and initial values from a log-uniform distribution. The parameters and initial values were varied two orders of magnitude above and below the initial guess (taken from previous models and literature evidence – see Supplementary File S1 “Parameters” and “Initial Values” spreadsheets). The total amount of phosphorylated STAT5 was calculated by summing together all forms of pSTAT5 and multiplying by two if the molecule included a STAT dimer with both STAT molecules phosphorylated.

We analyzed the features of the pSTAT5 concentration over time. The definitions of the characteristics of the pSTAT activation illustrated in Fig. 5A are as follows:

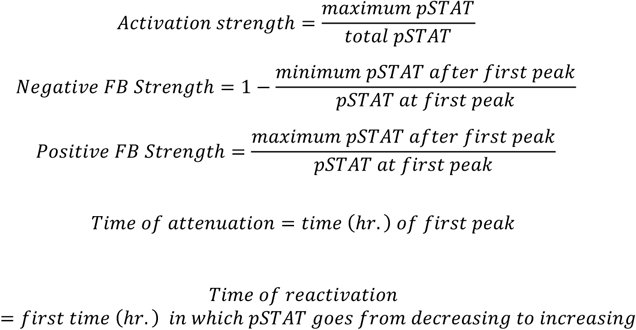

The number of peaks in total pSTAT5 was quantified using the Matlab *findpeaks* function, which returned the value of total pSTAT5 at local maxima as well as the time of the peak in hours. Thresholds for the *findpeaks* function were defined to have a minimum distance between peaks of 20 minutes and a minimum peak prominence of 0.1% to avoid identifying noise in the data as peaks (see MATLAB *findpeaks* documentation).

A detailed shape classification was performed based on the decision tree in Fig. S1 implemented through *if* statements in our MATLAB script. Parameter correlations were calculated in R using the *cor* function. The correlations shown in Fig. 5 are calculated using Monte Carlo simulations from the full model structure that included all three regulatory modules. Correlations with activation strength were calculated using all 100,000 simulations. Correlations with negative FB strength, positive FB strength, and time of attenuation could only be calculated for simulations that had a peak, *n* = 45,199. Correlations with the time of reactivation could only be calculated for simulations that had reactivation, *n* = 3,479.

### Model calibration

#### Sensitivity analysis

A total of 30 parameters were chosen to fit to the 31 experimental data points based on a global sensitivity analysis. We used the extended Fourier Analysis Sensitivity Test (eFAST) to determine which parameters significantly influence the model predictions.^30^ The eFAST method uses a variance decomposition method to determine the sensitivity of model outputs to model inputs. The first-order sensitivity *S*_*i*_ quantifies the fraction of variance in model output that is explained by the input variance in the parameter *i*.

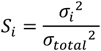

We calculated the first-order sensitivity of each kinetic parameter and non-zero initial value with the output being all species’ concentrations predicted by our model. We also estimated the total-order sensitivity *S*_*Ti*_ for each kinetic parameter and initial value. *S*_*Ti*_ is calculated as one minus the summed sensitivity index of complementary parameters *S*_*Ci*_ which is defined as all parameters except parameter *i*.

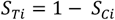

In order to determine which parameters to fit to experimental data, we compared the total-order sensitivity index for all kinetic parameters and initial values on the predicted model outputs: phosphorylated STAT5A (pSTATA), phosphorylated STAT5B (pSTATB), nuclear to cytoplasm ratio of STAT5A (STAT5A_n_ / STAT5A_c_), and the nuclear to cytoplasm ratio of STAT5A (STAT5B_n_ / STAT5B_c_). Although we calculated *S*_*Ti*_ for each parameter on all model outputs, we chose to focus on the effect of each parameter on those four model predictions because they are used in the objective function in model calibration (see below).

We took the mean *S*_*Ti*_ for each parameter or initial value over each of the four model outputs listed above at each timepoint for which we had experimental data from the literature. These sensitivity indices are included in the Supplementary File S1 on the sheet “Sensitivity results.” The parameters and initial values that had a mean *S*_*Ti*_ greater than that of the dummy variable, a factitious input which has no effect on model structure, were chosen as parameters to be fitted. In addition, the parameter *k30a*, which is the maximal rate of transcription of the PRLR receptor in response to STAT5 binding, was added to the parameter list because no kinetic parameter affecting the positive feedback module emerged from sensitivity analysis. In order to deconvolute the fact that the dimerization and shuttling rates of the different forms of STAT5 would likely be correlated, we defined the following multiplicative factors:

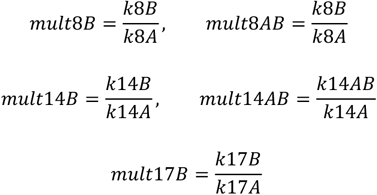

The parameters *k8A* and *k8B* describe the rate of homodimerization of STAT5A and STAT5B respectively while *k8AB* represents the rate of heterodimerization. The parameters *k14A, k14B*, and *k14AB* represent the rate of nuclear import of dimerized STAT5A dimers, STAT5B dimers, and heterodimers respectively. The parameters *k17A* and *k17B* represent the nuclear export rate of unphosphorylated STAT5A and STAT5B respectively.

#### Parameter estimation

Parameter fitting was performed by fitting the model simultaneously to all of the experimental data used for likelihood estimation. The amount of phosphorylated STAT5A and STAT5B at the 10 min., 30 min., 1 hr., 2 hr., 4 hr., and 6 hr. timepoints were quantified using Plot Digitizer (Java) from Brelje *et al*. Fig. 7.^11^ The nucleus to cytoplasm ratio of STAT5A and STAT5B at the 30 min., 1 hr., 1.5 hr., 3 hr., and 6 hr. timepoints and the 5 min., 15 min., 30 min., 1 hr., 2 hr., 3 hr., 4 hr., 5 hr., and 6 hr. timepoints respectively were quantified using Plot Digitizer from Brelje *et al*. Fig. 6 results for INS-1 cells.^11^ The fold change of the anti-apoptotic protein Bcl-xL in response for the timepoints 2, 4, 6, 8, 12, 18, and 24 hours were quantified using Plot Digitizer on Fujinaka *et al*. Fig. 7E.^19^ All experimental data from both papers was for INS-1 cells treated with 200 ng/mL of PRL.

A total of 25 independent fits were performed for each model structure using a Bayesian approach for likelihood estimation.^21,28,44^ Within each independent fit, 10,000 iterations on the parameter values were performed to effectively sample from the posterior distribution for each parameter value. The values from the iteration with the lowest error were taken as fitted parameters for each of the 25 independent fits. The formula for the sum of squared error is:

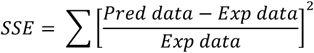

where *Pred data* and *Exp data* are vectors containing the model predictions and experimental measurements, respectively. Model predictions correspond to the same time points as the experimental data. For Table 1, AIC was calculated from median SSE values for each model structure as:

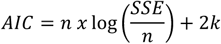

where *n* is the number of data points, *SSE* is the median error, and *k* is the number of parameters used to fit the model.

**Table 1:**
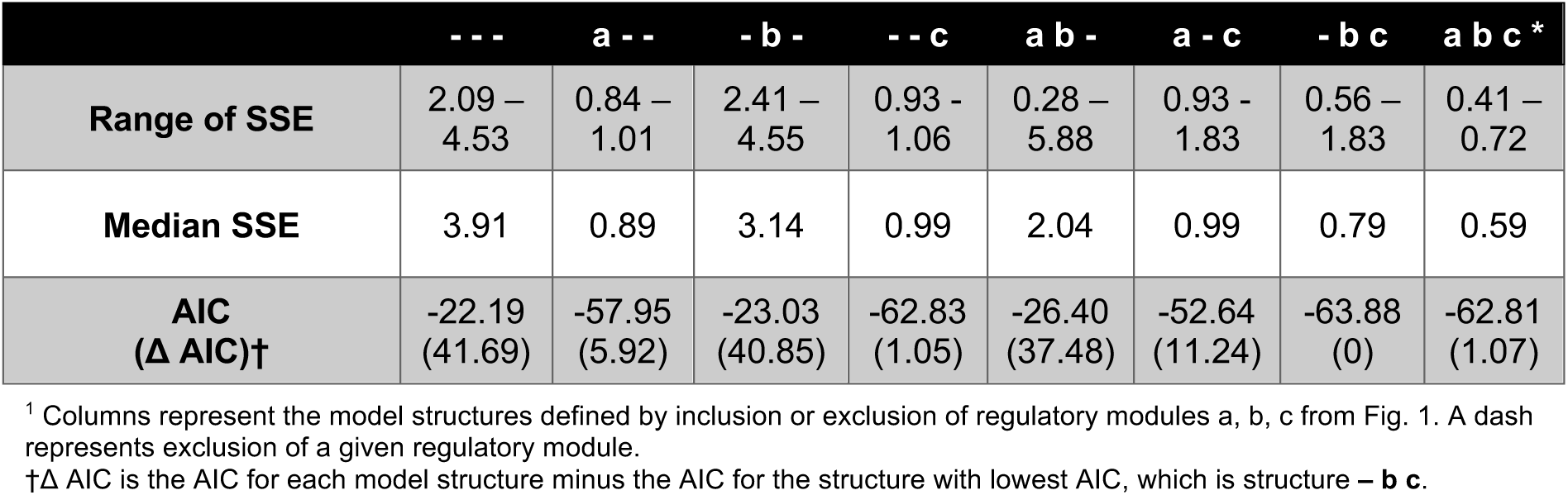
Comparison of Model Structures.

To display the results for Fig. 6 and Fig. S2-S8, the predicted time courses were simulated for each of the 25 independent fits per model structure. The mean and standard deviation of the model predictions were quantified and shown. For model validation (Fig. 7), the dose response data for rat primary beta cells was quantified using Plot Digitizer from Brelje *et al*. Fig. 13.^11^

## 3 RESULTS

### 3.1 Mechanistic model of JAK-STAT signaling in beta cells

A mechanistic model of JAK2-STAT5 signaling through the prolactin receptor was constructed based on known reactions from the literature.^8,26,34,39,43^ The model builds on the prior work of Yamada, *et al*. 2003 modeling control mechanisms in JAK-STAT signal transduction^48^ and Finley, *et al*. 2011, which analyzed IL-12 mediated JAK-STAT signaling in T cells.^16^

The mechanistic model includes a core network representing the canonical JAK-STAT signaling cascade, which includes 31 reactions and 24 molecular species (Fig. 1). Three regulatory modules are included or excluded from the network in order to consider their effect on STAT5 activation. These include: a) SOCS exerting negative feedback on STAT5 phosphorylation, b) receptor up-regulation due to transcriptional action of pSTAT5, and c) receptor internalization of the prolactin receptor induced by ligand binding. Including each regulatory module individually and in all possible combinations leads to eight model structures to explore. The full signaling network with all three regulatory modules included has 47 reactions and 32 molecular species. A full list of reactions is included in the supplementary material File S1.

### 3.2 Effect of different regulatory modules on qualitative shape of pSTAT5 activation

We defined eight model structures based on inclusion or exclusion of the three regulatory modules from Fig. 1 and ran Monte Carlo simulations for each structure to explore model predictions across a wide area of the parameter space. Here, we varied all parameter values (i.e., the kinetic reaction rates) and non-zero initial conditions (see Methods). This enables us to efficiently explore the parameter space and characterize the simulated dynamics that are generated. Each simulation was classified as “No peak”, “Single Peak”, or “Multiple Peaks” based on the predicted time course of STAT5 phosphorylation (Fig. 2A). Within the model structures with only one regulatory module included, the structure that included SOCS feedback was most likely to show multiple peaks in STAT5 phosphorylation (Fig. 2B). The structure that included receptor internalization was most likely to produce a single peak in STAT5 phosphorylation. Our simulations demonstrate that at least one regulatory module is required to produce multiple peaks in STAT5 phosphorylation (red values listed in Fig. 2B). Overall, the likelihood of randomly sampled parameter sets producing a time course of STAT5 phosphorylation with multiple peaks was very low for all model structures. Of the 8×10^5^ simulations we performed, only 0.2% (1,207) exhibited multiple peaks. This indicates that tight control of the reaction rates is necessary to achieve the right balance of activation and attenuation.

**Figure 2:**
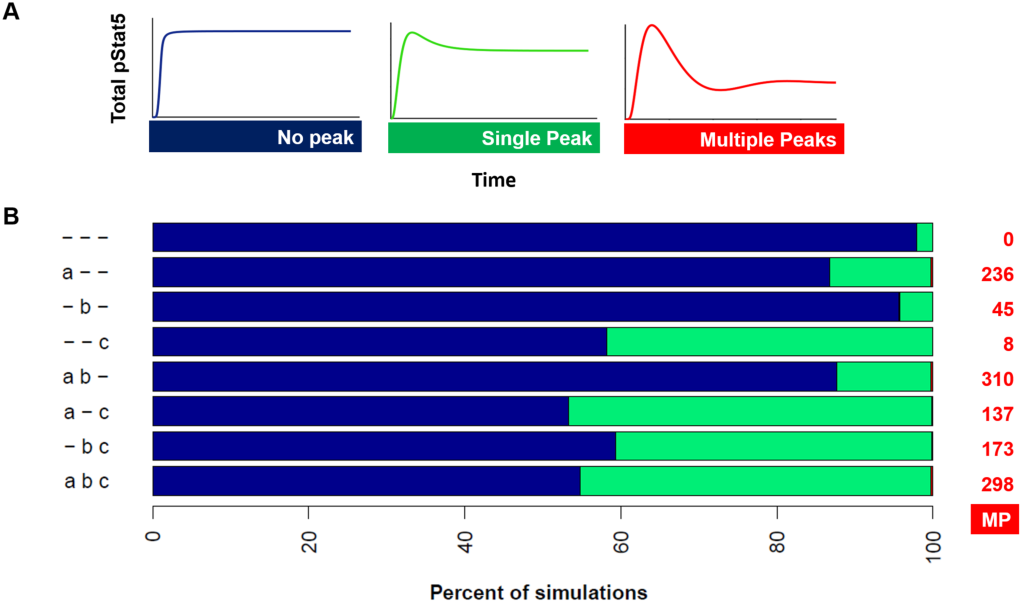
Ensemble Modeling Predicts the Number of Peaks in STAT5 Phosphorylation. **(A)** Simulated time courses were classified into three shapes based on the number of peaks in STAT5 phosphorylation over 6 hours of PRL stimulation. **(B)** Bar chart shows the percentage of Monte Carlo simulations from each model structure that were classified into each shape shown in panel A. Row labels correspond to the inclusion or exclusion of regulatory modules shown in Fig. 1. The data labels in red show the number of simulations that were classified as “Multiple Peaks” for each structure. *n* = 100,000 simulations per structure (800,000 total). MP, Multiple Peaks.

From the Monte Carlo simulations, model predictions that had multiple peaks in STAT5 phosphorylation showed wide variation in the magnitude and time course of phosphorylation. Therefore, we set out to define a more detailed classification to understand which model structures could give rise to STAT5 dynamics matching those observed in INS-1 cells, which show a defined profile for phosphorylated STAT5. Specifically, Brelje and colleagues showed that STAT5 reaches an initial peak at approximately 30 minutes following the initial stimulation, followed by attenuation between 1 – 3 hours, which reduces phosphorylation to below 70% of its initial peak. A second increase is observed after three hours, where phosphorylated STAT5 reaches or exceeds the initial level of phosphorylation.^10,11^ We first filtered the Monte Carlo simulations, retaining those that resulted in an appreciable level of STAT5 phosphorylation (at least 1% of the initial STAT5 becomes phosphorylated), assuming a minimum amount of phosphorylation is required to promote downstream signaling and cell response. We then defined eight qualitative shapes of STAT5 activation based on the number of peaks and the time at which the peaks occur. The decision tree used to classify predicted time courses is shown in Fig. S1.

This classification enabled a detailed characterization of the dynamics of phosphorylated STAT5. Almost one third of all simulations had no peak in STAT5 phosphorylation, showing the general shape of saturating kinetics (Fig. 3A). Almost 10% of simulations showed attenuation of the initial STAT5 activation leading to a single peak in pSTAT5 (Fig. 3B). A select few parameter sets (0.09%) produced multiple peaks in STAT5 phosphorylation that did not match the qualitative shape of experimental data, such as having more than one oscillation within 6 hours (Fig. 3C) or showing a smaller second peak characteristic of damped oscillation (Fig. 3D). Positive feedback can lead to unstable systems, and some simulations (0.05%) had an early peak in STAT5 phosphorylation followed by a large increase in phosphorylation due to strong positive feedback (Fig. 3E). Over 2,000 (0.25%) simulations had an initial peak followed by minimal attenuation before reactivation (Fig. 3F). These simulations are grouped into early, intermediate, and late simulations to preserve the qualitative shape when pooling simulations together. Another small group of simulations (0.02%) had multiple peaks in pSTAT5 but did not match the time course of the experimental data, either because the reactivation was too fast (< 3 hr) or the initial peak was too slow (> 1 hr) (Fig. 3G).

**Figure 3:**
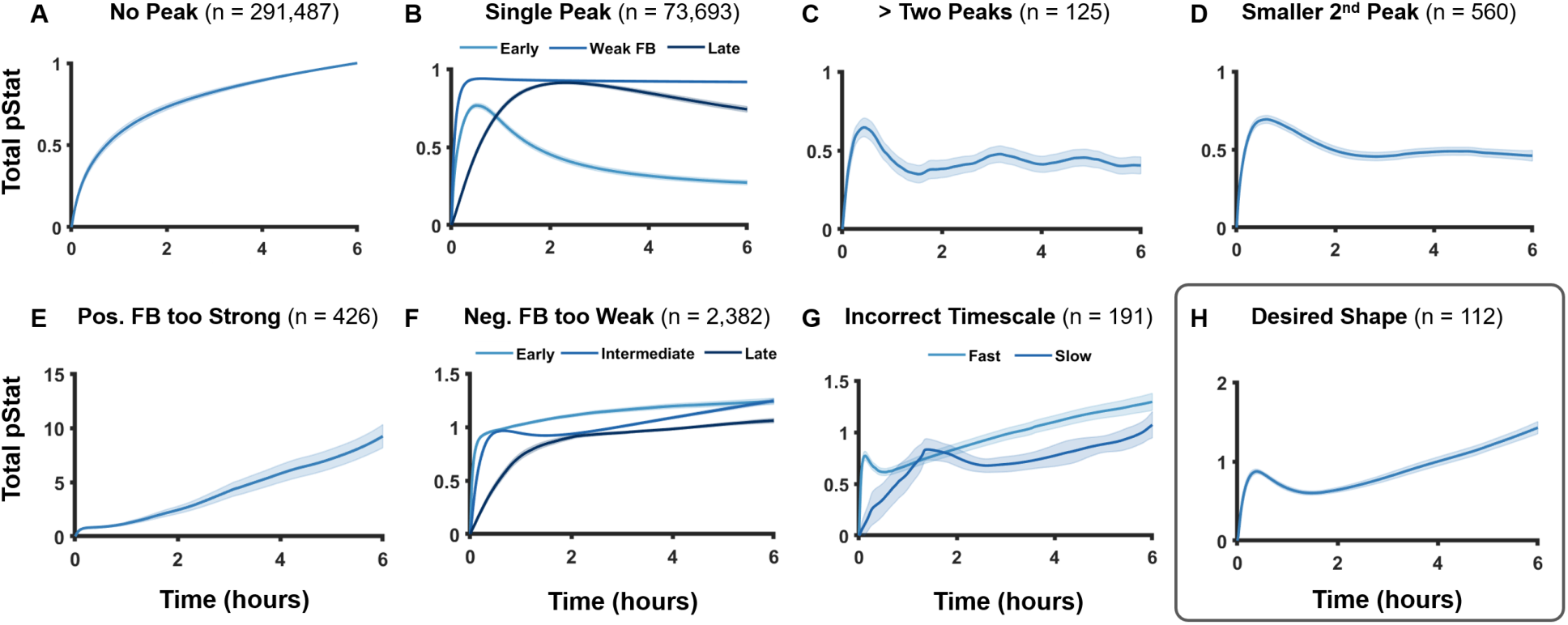
Classification of Simulations into Qualitative Shapes. Simulated time course of STAT5 phosphorylation for each shape shows the mean (solid line) and 95% confidence interval (shaded area) of all Monte Carlo simulations (800,000 total) classified into that shape. All shapes are mutually exclusive, that is, all simulations were uniquely assigned to one shape (see Figure S1 for decision tree). Simulations that did not reach a threshold level of 1% of STAT5 phosphorylated were labeled as “weak activation” and filtered out, n = 431,024.

Finally, a small number of simulations (112) matched the qualitative shape of the experimental data (Fig. 3H). Simulations classified as having this desired shape comprise just over 0.01% of the 800,000 total simulations, again pointing to the necessity of tightly controlled balance of positive feedback and negative feedback, both in terms of the strength and time-scale of feedback. The eight distinct model structures contributed differently to the fraction of simulations that match the desired shape (Fig. 4). Although SOCS inhibition was sufficient to cause multiple peaks in STAT5 phosphorylation (Fig. 2), SOCS inhibition alone was not sufficient to cause an early peak followed by prolonged activation (Fig. 4, row 2). Model structures that included receptor up-regulation, combined with either SOCS inhibition, receptor internalization, or both had the highest likelihood of matching the desired qualitative shape (Fig. 4, rows 5, 7, and 8).

**Figure 4:**
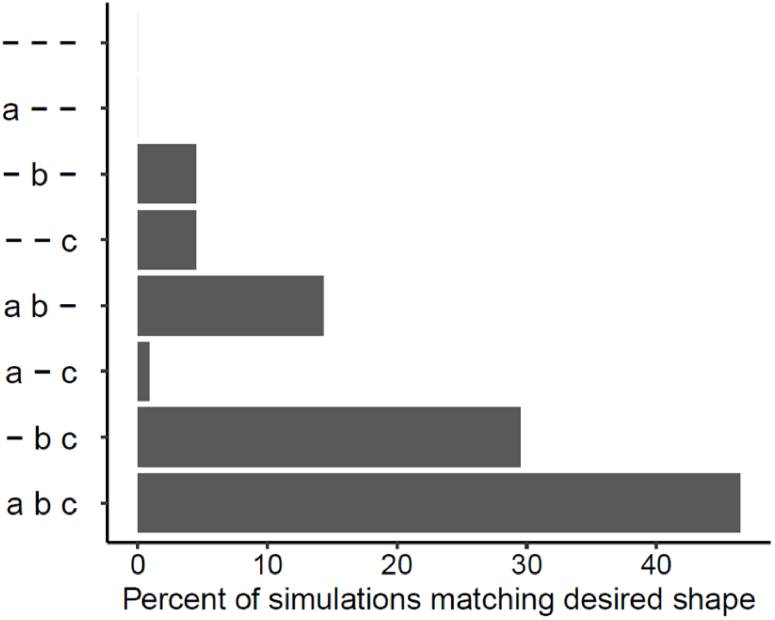
Breakdown of Simulations Matching Desired Shape by Structure. The y-axis shows the eight model structures defined by the inclusion or exclusion of the regulatory modules. Horizontal bars show the percentage contribution of each model structure to the 112 simulations which matched the desired shape shown in Fig. 3H.

### 3.3 Effect of parameter values on time course of STAT5 activation

Kinetic parameter values affect the strength of STAT5 activation, strength of feedback, and timescale of feedback. Several parameters from Monte Carlo simulations were strongly correlated with characteristics of the predicted time course of phosphorylated STAT5 (Fig. 5A). These characteristics include the activation strength, the strengths of the negative and positive feedback, and the times of attenuation and reactivation (see Methods section for more detail). The Pearson correlation coefficients for each influential parameter are shown in Fig. 5B-F, with the five parameters having the highest overall absolute value of the correlation coefficient labeled in each panel. The ratio of the ligand-bound receptor degradation rate to the unbound receptor degradation rate (*deg_ratio*) was highly correlated with all five defined characteristics of the pSTAT5 time course. As expected, higher values of *deg_ratio* decreased the activation strength (Fig. 5B), increased the strength of negative feedback (Fig. 5C), and decreased the strength of positive feedback (Fig. 5D). Increased values of *deg_ratio* also led to a faster timescale of attenuation (Fig. 5E) because the active receptor complex had a shorter half-life in the cell therefore less time to phosphorylate STAT5. Although high values of *deg_ratio* reduced the strength and timescale of initial STAT5 phosphorylation, they also lead to faster reactivation in simulations, which produced multiple peaks (Fig. 5F) since faster initial attenuation allowed for a faster rebound of signaling. The parameter *k2*, the ligand-receptor binding rate, had a similar effect as *deg_ratio* on the strength and timescale of feedback (Fig. 5C-F). However, it was positively correlated with the strength of activation. A faster rate of ligand binding leads to a stronger activation but stronger negative feedback due to increased internalization of ligand-bound receptors.

**Figure 5:**
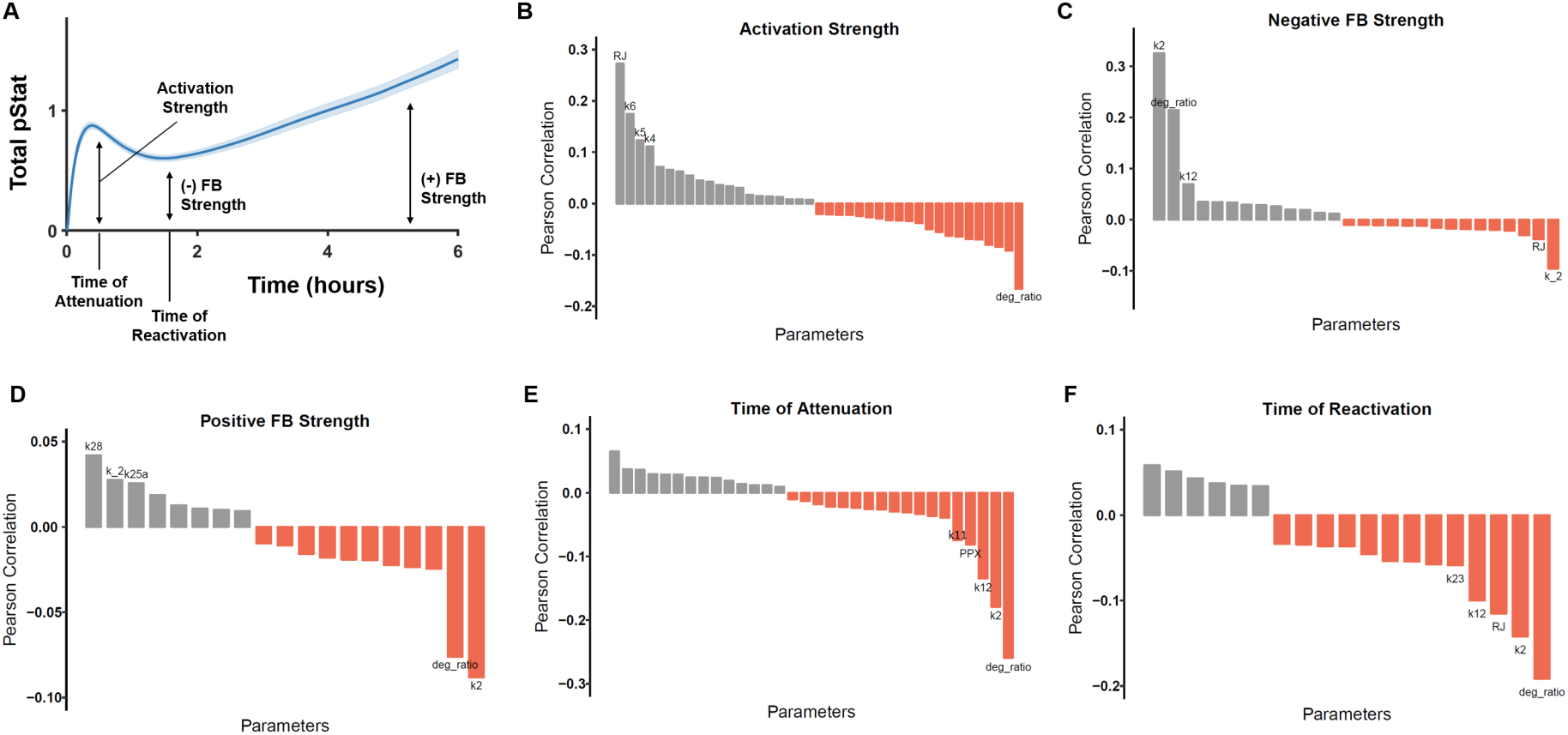
Parameters Correlated with STAT5 Phosphorylation. Pearson correlation between each kinetic parameter or initial value and five quantitative characteristics of the STAT5 phosphorylation time course. **(A)** Illustration of five characteristics. **(B)** Activation strength. **(C)** Negative feedback strength. (**D)** Positive feedback strength. **(E)** Time of attenuation. **(F)** Time of reactivation. Only parameters with statistically significant (*p* < 0 .05) correlations are shown in the waterfall plots. The five parameters most highly correlated with each characteristic are labeled. ***RJ***, initial value of PRLR:JAK2 complex; ***k6***, phosphorylation rate of STAT5; ***k5***, activation rate of JAK2; ***k4***, dimerization rate of PRLR:JAK2 complexes; ***deg_ratio***, ratio of degradation rate of ligand-bound receptor complexes to unbound complexes; ***k2***, ligand binding *on* rate; ***k12***, rate of dephosphorylation of pSTAT5 by cytoplasmic phosphatase; ***k_2***, ligand binding *off* rate; ***k28***, translation rate of PRLR mRNA; **k25a**, V_max_ for transcription of PRLR mRNA; ***k11***, binding rate of cytoplasmic phosphatase to pSTAT5; ***PPX***, initial value of cytoplasmic phosphatase; ***k23***, SOCS degradation rate.

Increased values of the initial concentration of the receptor:JAK2 complex increased the activation strength (Fig. 5B) and decreased the strength of negative feedback (Fig. 5C). Predictably, parameters that govern the rate of interactions critical to STAT5 activation (*k4, k5*, and *k6*, corresponding to the rate at which the ligand-bound receptor complex is activated, binds STAT5, and phosphorylates STAT5, respectively) were positively correlated with the activation strength (Fig. 5B). Additionally, increases in *k12*, the rate at which cytosolic phosphatase dephosphorylates STAT5, led to stronger negative feedback (Fig. 5C) and a faster timescale of attenuation (Fig. 5E). Overall, this analysis provides mechanistic insight into how specific biochemical reactions influence key features of STAT5 dynamics. Such results can guide experimental studies to modulate the signaling network to enhance STAT5 response.

### 3.4 Model calibration to STAT5 dynamics in INS-1 cells

The results presented thus far provide a detailed analysis of the model features that give rise to STAT5 dynamics that qualitatively agree with experimental data. We next aimed to produce a predictive model that quantitatively matches the data by calibrating the computational model to the experimental data. Specifically, we fit the model to measurements for the time course of phosphorylation of STAT5A and STAT5B^11^, translocation of STAT5A and STAT5B from the cytoplasm to the nucleus^11^, and the fold change in protein level of the response protein Bcl-xL.^19^

Fitting all eight possible model structures to the experimental data reveals the relative importance of regulatory modules (Table 1). The model structure that included all three regulatory modules from Fig. 1 had the lowest median Sum of Squared Errors (SSE) and tightest range of SSE. Predictably, model structures which did not include receptor up-regulation (module b) could not achieve a second peak in STAT5 activation that was higher than the first (Fig. S2, S3, S5, S7). This result matches the results presented above in Results section 2.2. The model structure with just SOCS feedback (regulatory module a) showed damped oscillations, in agreement with prior computational modeling of JAK-STAT regulation^48^ (Fig. S3). The model structure with just receptor internalization (module c) showed a strong attenuation of the initial peak but no reactivation (Fig. S5) and the model structure that only included receptor up-regulation (regulatory module b) had very high SSE values because negative feedback was not strong enough to attenuate the initial peak in STAT5 activation (Fig. S5). Interestingly, the model structure that included only SOCS feedback and receptor up-regulation (modules a and b) had the lowest minimum SSE but the widest variation in SSE (Table 1). The high variation in fitted error value reflects the instability of a system with strong negative and positive feedback. Kinetic parameters must be tightly controlled for a system like this to achieve an oscillatory response.

In addition to using the SSE to evaluate the model fits, we also use the Akaike Information Criterion (AIC), which allows for comparison of model structures with different number of fitted parameters, penalizing structures that have more parameters.^37,40^ A lower value of AIC indicates a better fit. The model structure without SOCS negative feedback had the lowest AIC. This structure fit the data similarly well as the full model, albeit with a wider variability in SSE. In summary, by studying the various model structures containing different sets of regulatory mechanisms, we are able to more fully understand the mechanisms required to match experimental data.

We analyzed the estimated parameter values and predictions for the model that included all three regulatory modules, as that model structure produced the lowest median SSE and provided consistent parameter estimates. Model predictions for this full model are shown in Fig. 6, illustrating that this structure effectively captured the phosphorylation dynamics (Fig. 6A-B) and the nuclear import (Fig. 6C) of both STAT5A and STAT5B on the six-hour timescale. The dynamics of these two species share a similar qualitative shape because phosphorylation is necessary for shuttling of STAT5 to the nucleus.

**Figure 6:**
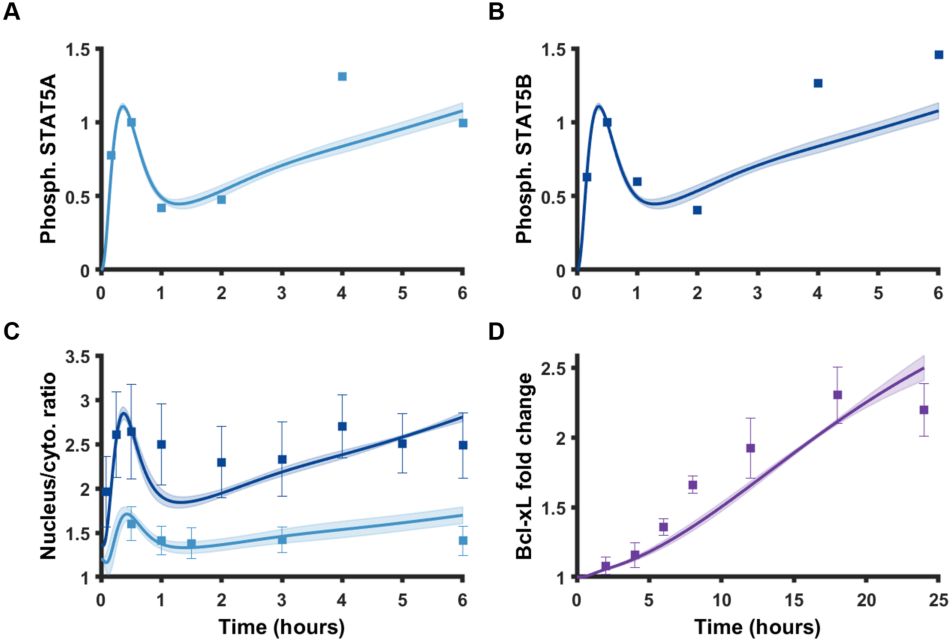
Model calibration. Model predictions for **(A)** Phosphorylated STAT5A, normalized to the 30 minute time point, **(B)** Phosphorylated STAT5B, normalized to the 30-minute time point, **(C)** Ratio of nuclear to cytosolic phosphorylated STAT5, and **(D)** Fold change of Bcl-xL. Lines show mean value of model predictions with shading indicating the standard deviation for 25 independent fits. Squares show experimental data points from Brelje *et al*. for panels A, B, and C or from Fujinaka *et al*. for panel D. Error bars are included for experimental data points that had error bars shown in the previously published work. All experimental data are for INS-1 cells treated with PRL at 200 ng/mL. Thirty parameters were fit simultaneously to the five data sets using a Bayesian likelihood estimation approach. *Dark blue*, STAT5A; *light blue*, STAT5B; *purple*, Bcl-xL.

Interestingly, although STAT5A and STAT5B show a similar pattern in phosphorylation, they differ in the amount that is translocated into the nucleus.^10,11^ The model accounts for separate STAT5A and STAT5B species and allows for homo- and hetero-dimerization with separate rate constants for dimerization, import of phosphorylated dimers into the nucleus, and export of dephosphorylated STATs from the nucleus. The fitted model predicts that STAT5B homodimers and STAT5AB heterodimers dimerize nearly three times faster than STAT5A homodimers, with ratios of 2.89 ± 1.15 and 2.78 ± 1.37, respectively as compared to the STAT5A dimerization rate. The model also predicts the nuclear import rate to be faster for STAT5B homodimers and STAT5AB heterodimers than STAT5A homodimers, with ratios of 2.80 ± 1.12 and 2.40 ± 1.04, respectively. The faster dimerization rate and nuclear import rate explains how more STAT5B localized to the nucleus compared to STAT5A.

In addition to predicting the upstream dynamics, the model also captures how a single oscillation in STAT5 activation on the six hour timescale can lead to a smooth increase in the concentration of a response protein on a longer time-scale (Fig. 6D). We note that in both the full model and the structure without SOCS feedback, the dimerization rate and import rate of STAT5B and STAT5AB were faster than STAT5A, showing that this conclusion holds across multiple model structures.

Taken as a whole, the fitting results suggest that multiple feedback mechanisms could explain the observed time courses in STAT5 phosphorylation, nuclear translocation, and protein response. However, receptor upregulation is required, and it must be combined with at least one of the other regulatory mechanisms (SOCS negative feedback or receptor internalization). The calibrated model containing all three regulatory modules produces the best fit to the data and generates consistent parameter estimates (Fig. S10).

### 3.5 Validation of model predictions with data from rat primary beta cells

We aimed to validate the calibrated model containing all three regulatory modules using separate experimental observations. The dose response curve for the ratio of STAT5B in the nucleus compared to the cytoplasm predicted by the computational model using fitted parameter values qualitatively matches what is observed for rat primary beta cells *in vitro* after 30 minutes of stimulation with PRL^11^ (Fig. 7A). Importantly, both the experiments and model predictions show a biphasic response, in which the STAT5B level increases with increasing stimulation before decreasing. However, there is a difference in the time at which the maximum response predicted by the model matches the experimental measurements. Specifically, the model predicts an increase in STAT5B translocation at the 30-minute timepoint (Fig. 7A, light blue bars) with increasing hormone concentration, with the maximal response occurring at a dose of 500 ng/mL. In comparison, the peak response occurs at the 1,000 ng/mL dose in the experimental data (Fig. 7A, grey bars). Given that the model produces the full time course of STAT5 levels, we can investigate why there is this difference between model and experiments. The model predicts that the attenuation of the initial STAT5 activation occurs more rapidly for higher doses of PRL such that attenuation has already reduced STAT5 levels by the 30-minute timepoint (Fig. S9). We further analyzed the predicted STAT5 time course and found that considering the 18-minute timepoint (Fig. 7A, dark blue bars) gives a maximum dose response at 1,000 ng/mL of PRL, which matches the experimental data. This difference in the timing may be due to having calibrated the model using *in vitro* measurements, while the model validation data come from *in vivo* experiments. Although one study found high concurrence between *in vitro* and *in vivo* enzyme catalytic rates^13^, another recent study showed lower catalytic efficiency of a specific enzyme i*n vivo* versus *in vitro* ^50^.

**Figure 7:**
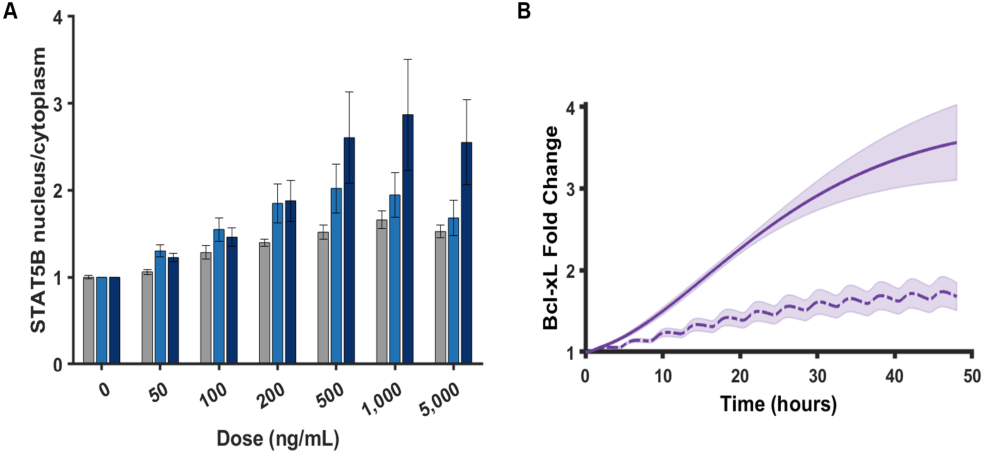
Model validation. **(A)** Model predicted dose response data for 30-minute timepoint (light blue) and 18-minute timepoint (dark blue), compared to experimental data from Brelje *et al*. treating rat primary beta cells with PRL shown in grey. Values are normalized to the amount of STAT5B in the nucleus with no PRL stimulation (0 ng/mL dose). Height of bars for model predictions show mean of 25 independent parameter sets with error bars indicating standard deviation. **(B)** The fold change in the concentration of the response protein Bcl-xL when treated with 200 ng/mL of PRL continuously for 48 hours (solid line) or pulsed with 1 hour of hormone every 4 hours (dashed line) to simulate the conditions tested by Brelje, *et al*. in rat primary beta cells. The line shows the mean model prediction from the 25 sets of fitted parameters with shading ± one standard deviation.

We also compared the model predictions to experimental data for Bcl-xL. Experiments conducted in rat primary beta cells showed that cell proliferation (as measured by Brdu-labeled nuclei) was about twice as high under 48 hours of continuous PRL treatment versus 48 hours of pulsed treatment. In the case of pulsed treatment, the researchers alternated between 1 hour of stimulus and 3 hours of wash.^10^ Using the fold change of the anti-apoptotic protein Bcl-xL as a proxy for cell proliferation, our fitted model predicts that the fold change of Bcl-xL is approximately two-times higher for continuous treatment versus pulsed (Fig. 7B).

Overall, considering both of these data sets, the model is validated by experimental observations. This establishes confidence that the model can generate reliable predictions and quantitative insight into JAK2-STAT5 signaling mediated in pancreatic beta cells.

### 3.6 Perturbing the fitted, validated model

After calibrating and validating the model, we examined the influence of varying individual parameters and initial values on model predictions. We varied each parameter or initial value that was determined by ensemble modeling to have a large impact on one of the various aspects of STAT5 activation (Fig. 5B-F) individually within two orders of magnitude of the fitted values. In general, changing individual parameter and initial values could alter both the strength of activation and the feedback dynamics, suggesting that the feedback system can be modulated. The impact of altering the kinetic parameters *k2* and *k12* as well as the initial values of *RJ* and *PPX* on the nuclear import of STAT5A and STAT5B as well as the Bcl-xL fold change are shown in Fig. S12.

We chose to investigate in detail two of the highest ranking influential kinetic parameters and initial values, based on their ability to strongly modulate multiple aspects of STAT5 activation. We quantified the initial peak in STAT5B nucleus to cytoplasm ratio which represents the strength of activation of the system. When varying the PRL ligand binding rate (*k2*) and the cytosolic phosphatase dephosphorylation rate (*k12*) two orders of magnitude, we found that the activation of STAT5 was more sensitive to changes in the ligand binding rate, as indicated by the increase in activation along the y-axis (Fig. 8A). The phosphatase did modulate activation, with higher values of *k12* leading to lower activation, but the effect is less pronounced than that of *k2*. A similar result was obtained when varying the initial value of the receptor:JAK complex (*RJ*) and the cytosolic phosphatase (*PPX*). The activation was increased greatly when the initial value *RJ* approached ten times its fitted value (Fig. 8B). Higher initial values of *PPX* decreased the strength of initial value, but again, this effect is less pronounced than modulating signaling at the receptor level. Based on these results, we conclude that targeting parameters upstream in the signaling network has a larger impact on the activation of STAT5, as compared to altering kinetic parameters and species further down the signaling cascade.

**Figure 8:**
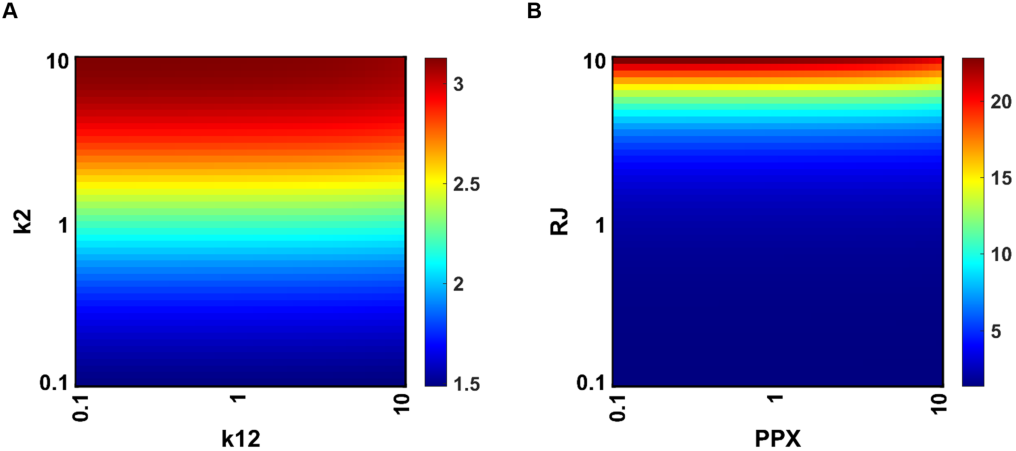
Model perturbations. **(A)** The effect of varying the initial ligand-binding rate *k2* and the cytosolic phosphatase dephosphorylation rate *k12* between 0.1-fold and 10-fold of the fitted parameter values. **(B)** Varying the initial values of the receptor-JAK2 complex *RJ* and the cytosolic phosphatase *PPX* between 0.1-fold and 10-fold of the fitted values. Coloring of the heat map indicates the initial peak in the STAT5B cytoplasm to nucleus ratio, averaged over the 25 independent fitted parameter sets for the full model structure including all regulatory modules.

## 4 DISCUSSION

Our mechanistic model of JAK-STAT signaling in pancreatic beta cells captures key dynamics of STAT5 activation via phosphorylation by the PRLR-JAK2 complex, followed by import of phosphorylated STAT5 dimers into the nucleus. The model differentiates between STAT5A and STAT5B and identifies the kinetic rate parameters that are able to explain experimentally observed differences in the amount of the STAT5A and STAT5B entering the nucleus under prolactin stimulation.^10,11^ Specifically, the model shows that the rates of dimerization and translocation can account for the experimental measurements. This mechanistic insight is relevant, as it has been hypothesized that this differential expression of STAT5A and STAT5B in the nuclear and cytosolic forms may be a form of tissue-specific regulation of JAK-STAT signaling^2^, arising from their differential affinity for STAT5 target genes.^6^ The model simultaneously predicts experimental data for upstream activation of STAT5 and fold change of the response protein Bcl-xL to the same hormonal stimulus. In addition, the validity of the model predictions of STAT5 nuclear import was explored by simulating dose response curves and comparing continuous versus pulsed simulations. In both cases, the model recapitulated data that was not used to train model parameters.

Our model validation is significant for two reasons. We show that the model can qualitatively and quantitatively match experimental data not used in model fitting. This demonstrates the predictive capability of the model. In addition, we establish that although the model was calibrated using data from INS-1 cells, it can reproduce observations obtained using primary pancreatic beta cells. This is a particularly important point since INS-1 cells, while used as a model of primary beta cells, exhibit quantitative differences in their metabolism^42^ and insulin secretion in response to glucose^32^, as compared to healthy beta cells. Thus, the model predictions are valid and reliable for a range of settings.

The model includes reactions known to drive JAK-STAT signaling in pancreatic beta cells. There are a multitude of feedback modules affecting the signal transduction pathway^34,39^, and we chose to explicitly explore the role of different feedback modules on the activation of STAT5 through ensemble modeling. We hypothesized that positive feedback through STAT5-induced receptor up-regulation could explain the reactivation of STAT5 in INS-1 cells to a magnitude greater than the initial activation.^10,11^ Classifying Monte Carlo simulated time courses by their qualitative shapes revealed that model structures with both receptor up-regulation and an inhibitory module (whether that be SOCS feedback of receptor internalization) were most likely to show reactivation of STAT5, matching the shape of the experimental data. Quantifying the impact of different parameter values on the time course of STAT5 activation helped us define which parameters drive the dynamics. An increased ligand-bound receptor degradation rate, for example, decreased the strength of activation and timescale of feedback while increasing the negative feedback strength. We followed up on the most influential parameters from ensemble modeling by varying them within the fitted, validated model. Our simulations predict that modulating signaling at the receptor level produces larger increases in STAT5 activation than altering the effect of an individual feedback mechanism (cytosolic phosphatase). This information is relevant for researchers aiming to enhance beta cell mass through activation of the JAK-STAT pathway. Ultimately, we found that multiple model structures could fit the data well (Table 1), but there were emergent properties that were consistent across model structures, such as a faster rate of STAT5B dimerization and nuclear import, as compared to STAT5A.

Our model motivates new experiments that can better elucidate the role of regulatory elements in JAK-STAT signaling. We are drawn to the work of Apgar *et al*. on stimulus design for model selection and validation^3^ as a logical next step of our work. Based on our findings, multiple possible inhibitory mechanisms could explain the observed time course of STAT5 phosphorylation. By designing a time course stimulus of PRL on INS-1 cells that aims to discriminate between these different model structures, one could experimentally test which mechanism is most likely to occur within the cell. This in-depth exploration of signal transduction would benefit pre-clinical researchers trying to design a therapy aimed at increasing beta cell mass in model organisms of diabetes.

Taken as a whole, our work points to the importance of regulatory modules in JAK-STAT signaling within pancreatic beta cells. Our model predicts that positive feedback combined with inhibition, be that through negative feedback or enhanced degradation rate, can drive a single oscillation in STAT5 phosphorylation within six hours, followed by a second peak that is higher than the first. Based on the rarity of this behavior occurring within the wide parameter space sampled, we contend that the kinetic rate parameters within the cell must be well constrained to balance positive and negative feedback and achieve this behavior. In line with this hypothesis, the kinetic parameters predicted by our model when fitting to experimental data were tightly constrained (Fig. S10).

Excitingly, the mechanistic insight as to the detailed effects of the regulatory modules provides quantitative information needed to identify strategies to increase beta cell mass. The ability to increase the beta cell mass *in vivo* could be a powerful new therapy for the treatment of diabetes.^14^ Hormonal stimulus seeks to recapitulate the islet adaptation to pregnancy^5^ and has already achieved beta cell proliferation in rodent models^9^ in both female and male rodents.^22^ Despite these advances, potential therapies have failed to realize the same gains in beta cell proliferation in humans^8,12,26,43^, pointing to a need for better understanding of regulatory mechanisms through the PRLR-JAK-STAT pathway.^12^ Here, we provide evidence that feedback modules play a key role in regulation of JAK-STAT signaling within a computational model relevant to the pancreatic beta cell. We also show that modulating upstream parameters such as the ligand binding rate and the initial value of receptor complexes can increase PRL-mediated STAT5 activation. We acknowledge the dependence of our model predictions on the accuracy of the model structure, and therefore explored several potential structures through ensemble modeling. The inclusion and exclusion of different regulatory modules gives insight into their relative importance and helps us understand the important predicted behaviors that emerge across multiple model structures.

## ACKNOWLEDGEMENTS

The authors would like to thank Dr. Mahua Roy, Dr. Jennifer Rohrs, Qianhui (Jess) Wu, Min Song, Sahak Makaryan, Ding Li, Colin Cess, and Patrick Gelbach for their suggestions and support. Sahak Makaryan provided valuable guidance regarding likelihood estimation parameter fitting. The work was partially supported by the USC Provost’s Undergraduate Research Fellowship and the Undergraduate Research Associates Program (URAP) awarded to R.D. Mortlock and USC Diabetes & Obesity Research Institute Pilot Funding Grant to

S.K. Georgia and S.D. Finley

## SUPPORTING INFORMATION

**File S1.** Model reactions, species, and parameter values.

**File S2.** Supplementary figures.

