## Supplementary material for "Dynamic regulation of JAK-STAT signaling through the prolactin receptor predicted by computational modeling": File S2

Figure S1: Decision Tree for Shape Classification

All 800,000 Monte Carlo simulations were grouped into eight mutually exclusive shapes based on the prediction for total amount of pSTAT5, which is abbreviated as pStat in the figure. The abbreviation "pks" describes the vector of local maxima in pStat model predictions returned by Matlab's *findpeaks* function. The abbreviation "loc" describes the time at which the local maxima occur. Similarly, the abbreviation "vl_loc" describes the time values corresponding to local minima in the pStat predictions. The decision tree reads from left to right along the gold line, meaning that once a decision is made that brings a shape "off of the gold line", subsequent decisions are not made on that simulation. Thus, the simulations matching the desired shape are those which have not been filtered out by any of the logical statements shown in the decision tree.

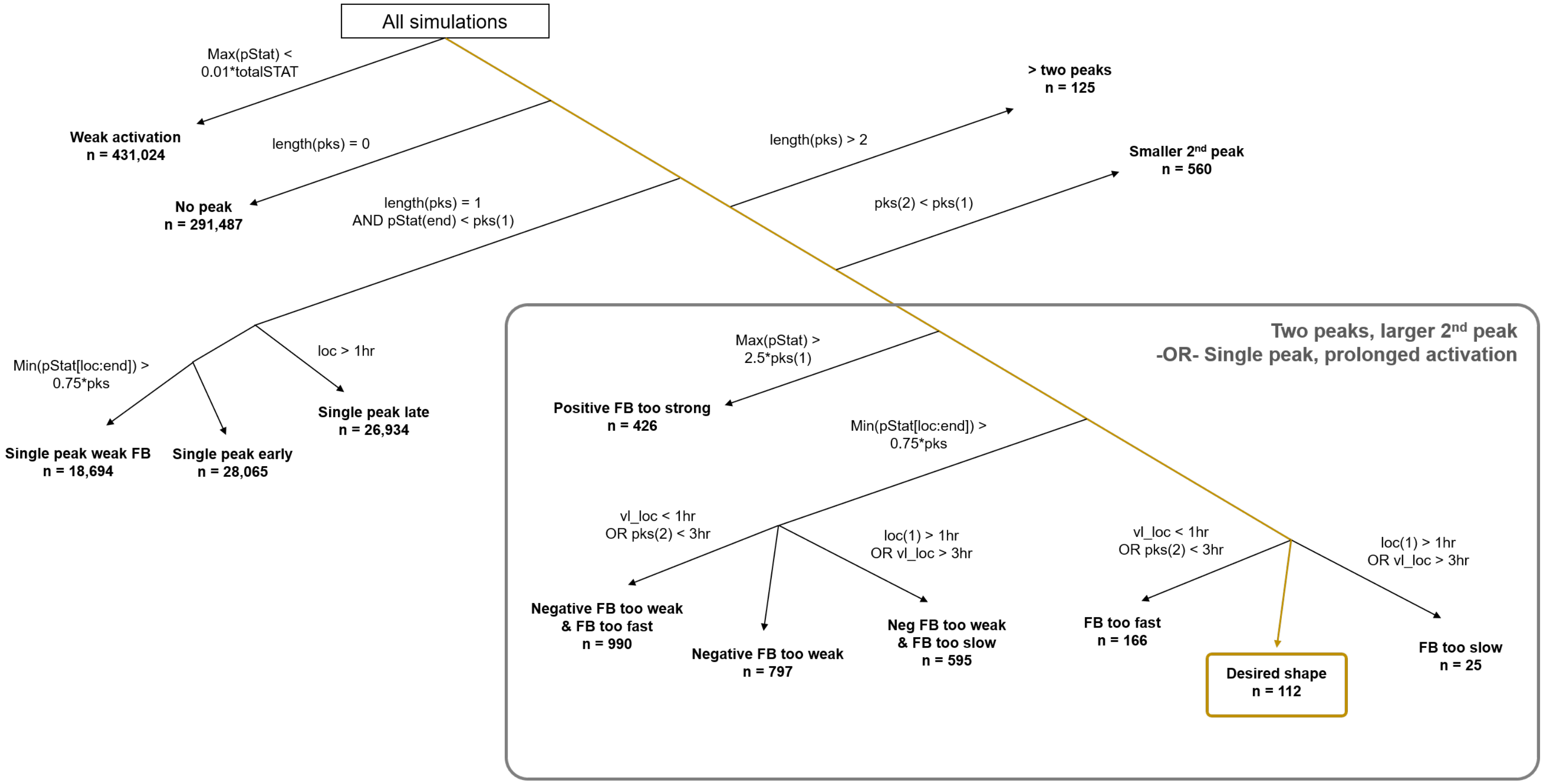

Figures S2-S8: Model Calibration for All Model Structures

Lines show mean value of model predictions with shading showing the standard deviation for 25 independent fits for various model structures. The letters in the top left correspond to which of the regulatory modules from Fig. 1 are included. Squares show experimental data points from Brelje *et al.* for panels A, B, and C or from Fujinaka *et al.* for panel D. Error bars are included for experimental data points that had error bars shown in the previously published work. All experimental data is for INS-1 cells treated with PRL at 200 ng/mL. Thirty parameters were fit simultaneously to the five data sets using a Bayesian likelihood estimation approach. *Dark blue,* STAT5A; *light blue,* STAT5B; *purple,* Bcl-xL

**Figure S2**

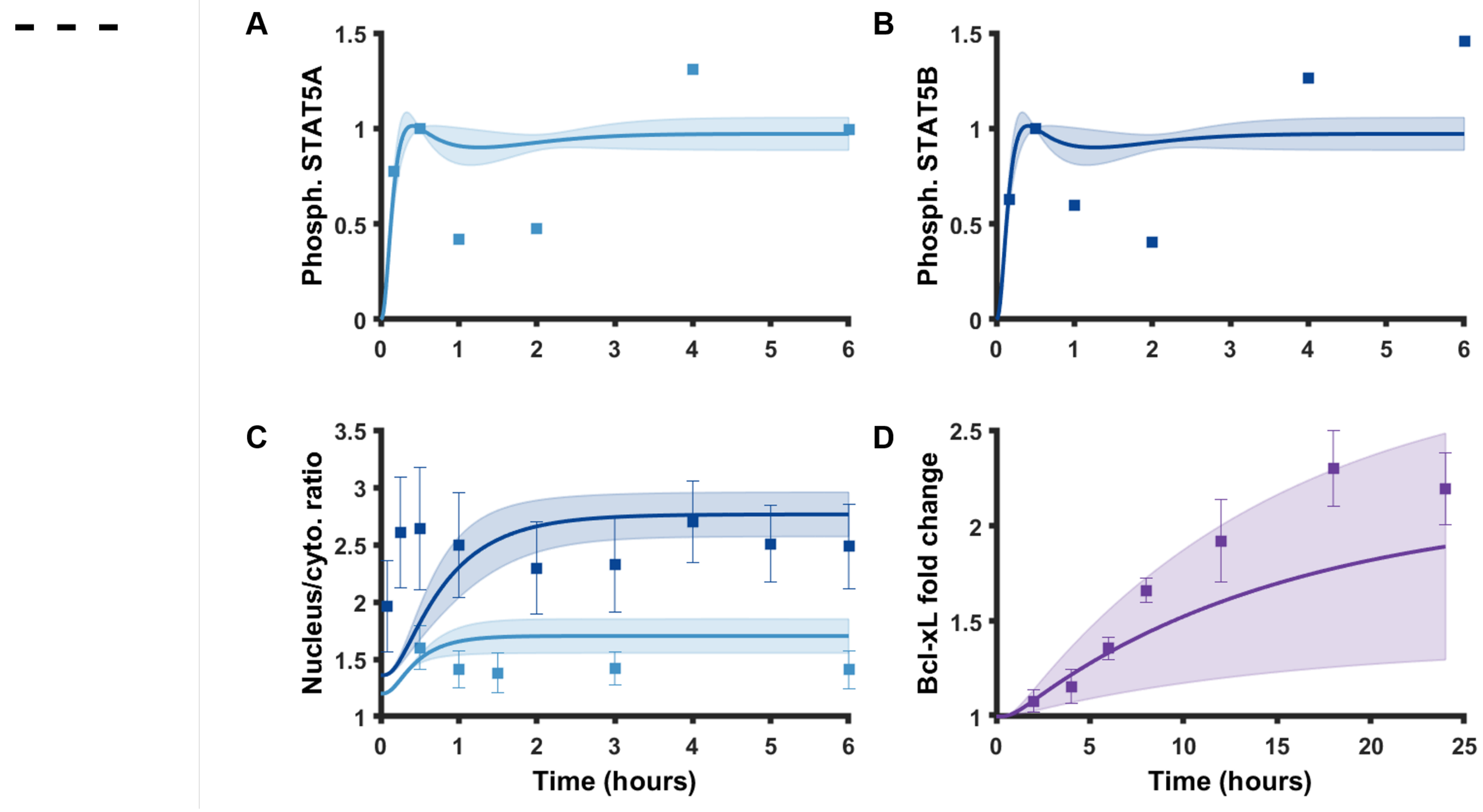

**Figure S3**

a - -

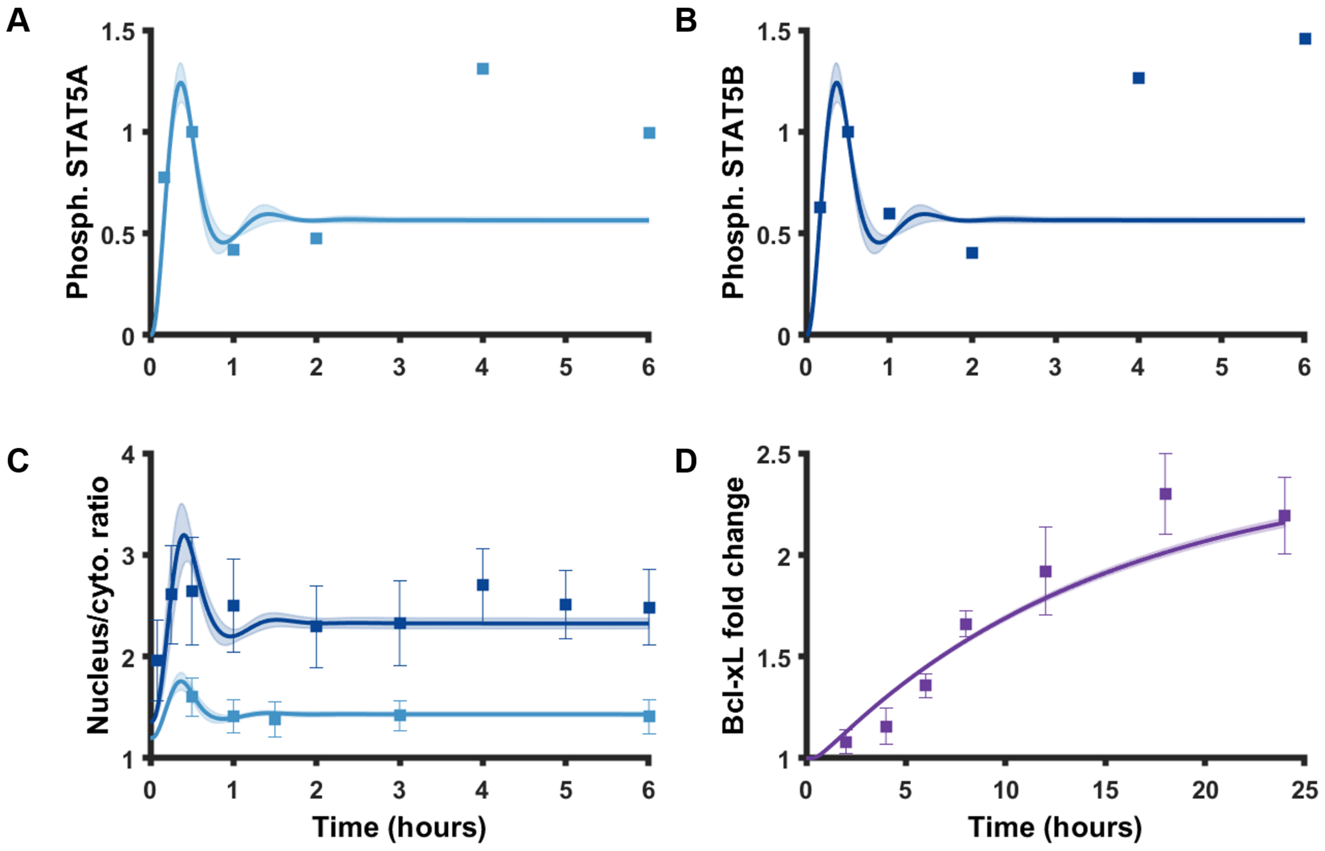

A
B
C
D
Phosph. STAT5A
Phosph. STAT5B
Nucleus/cyto. ratio
Bcl-xL fold change
Time (hours)
Time (hours)
1.5
1
0.5
0
0 1 2 3 4 5 6
4
3
2
1
2.5
2
1.5
1
0 5 10 15 20 25

**Figure S4**

- b -

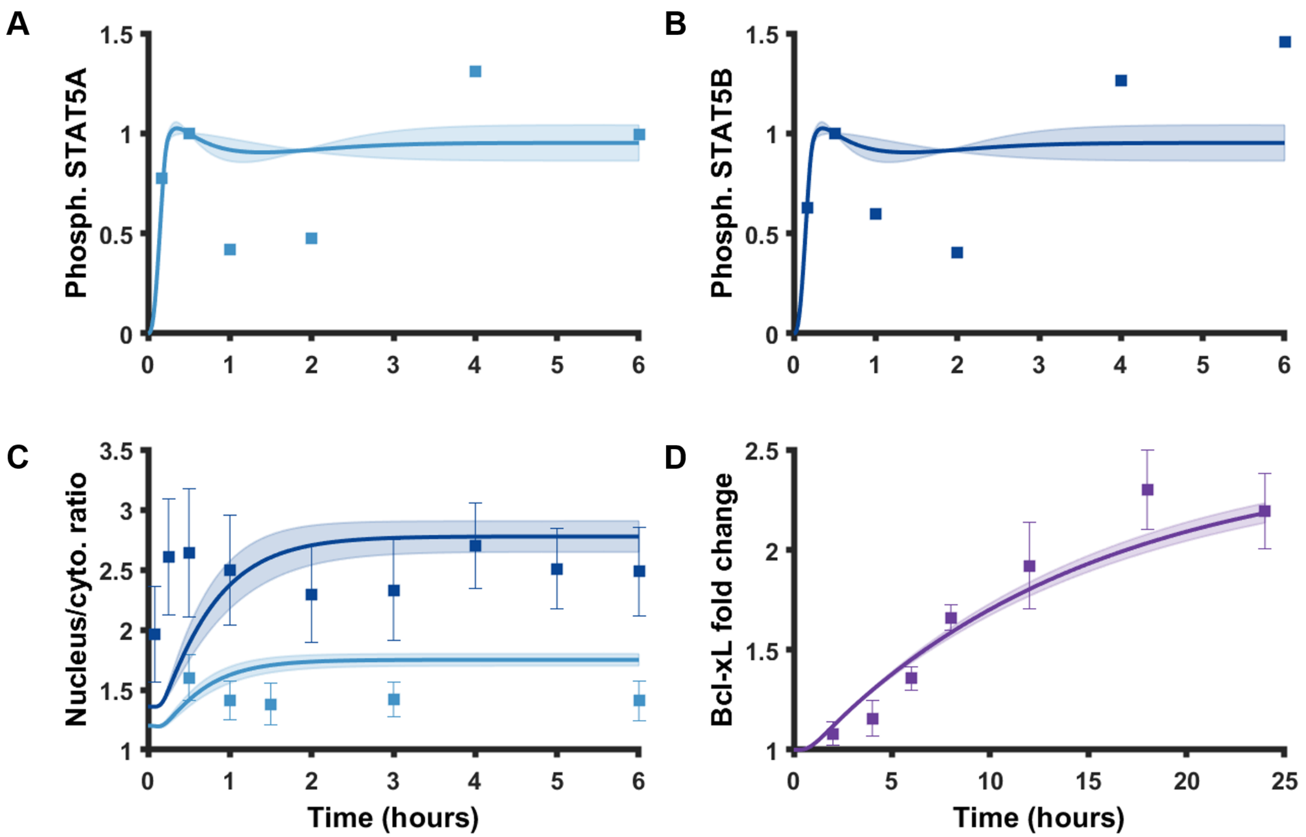

A
Phosph. STAT5A
1.5
1
0.5
0
0
1
2
3
4
5
6
B
Phosph. STAT5B
C
Nucleus/cyto. ratio
3.5
3
2.5
2
1.5
Time (hours)
D
Bcl-xL fold change
10
15
20
25

**Figure S5**

**- - C**

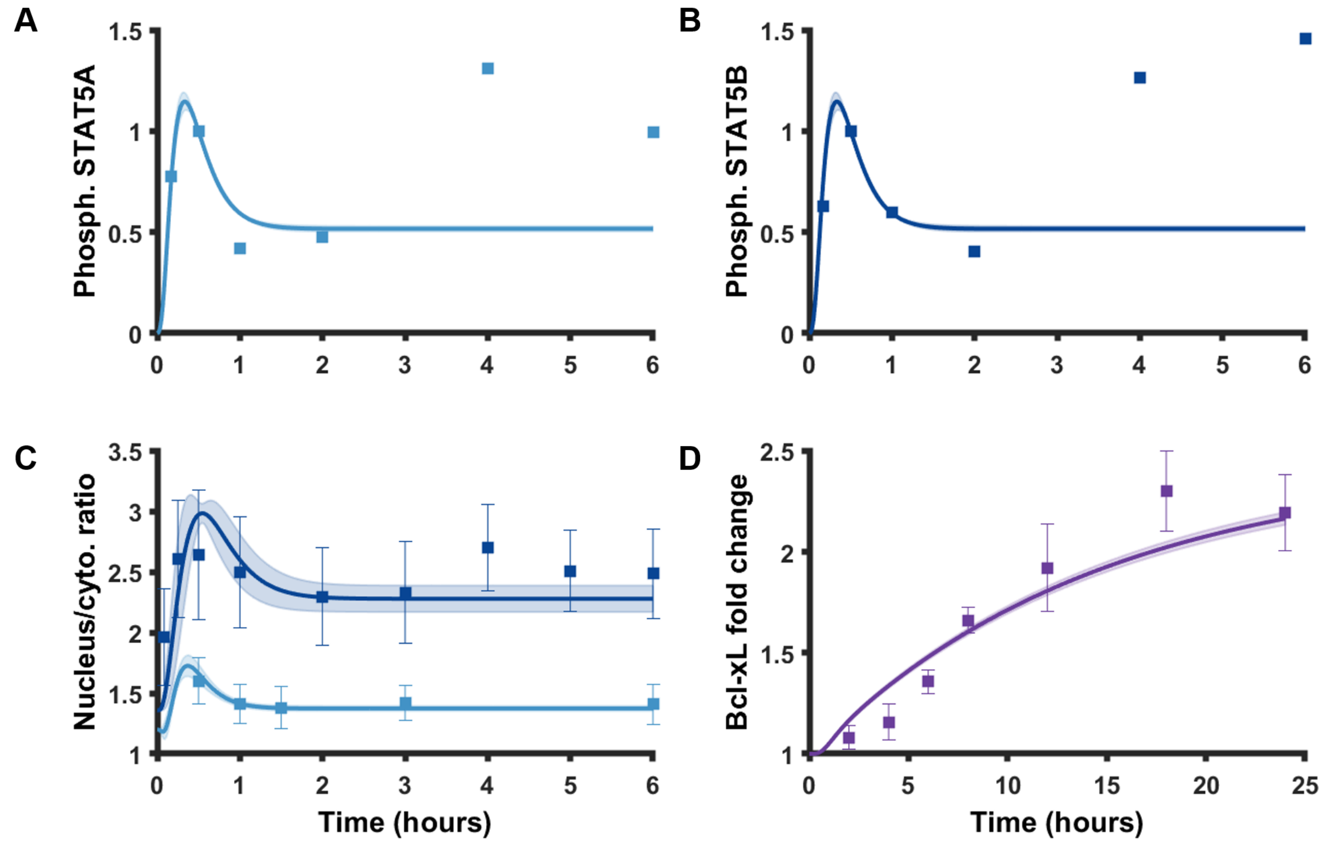
A
B
C
D
Phosph. STAT5A
Phosph. STAT5B
Nucleus/cyto. ratio
Bcl-xL fold change
Time (hours)
Time (hours)

**Figure S6**

a b -

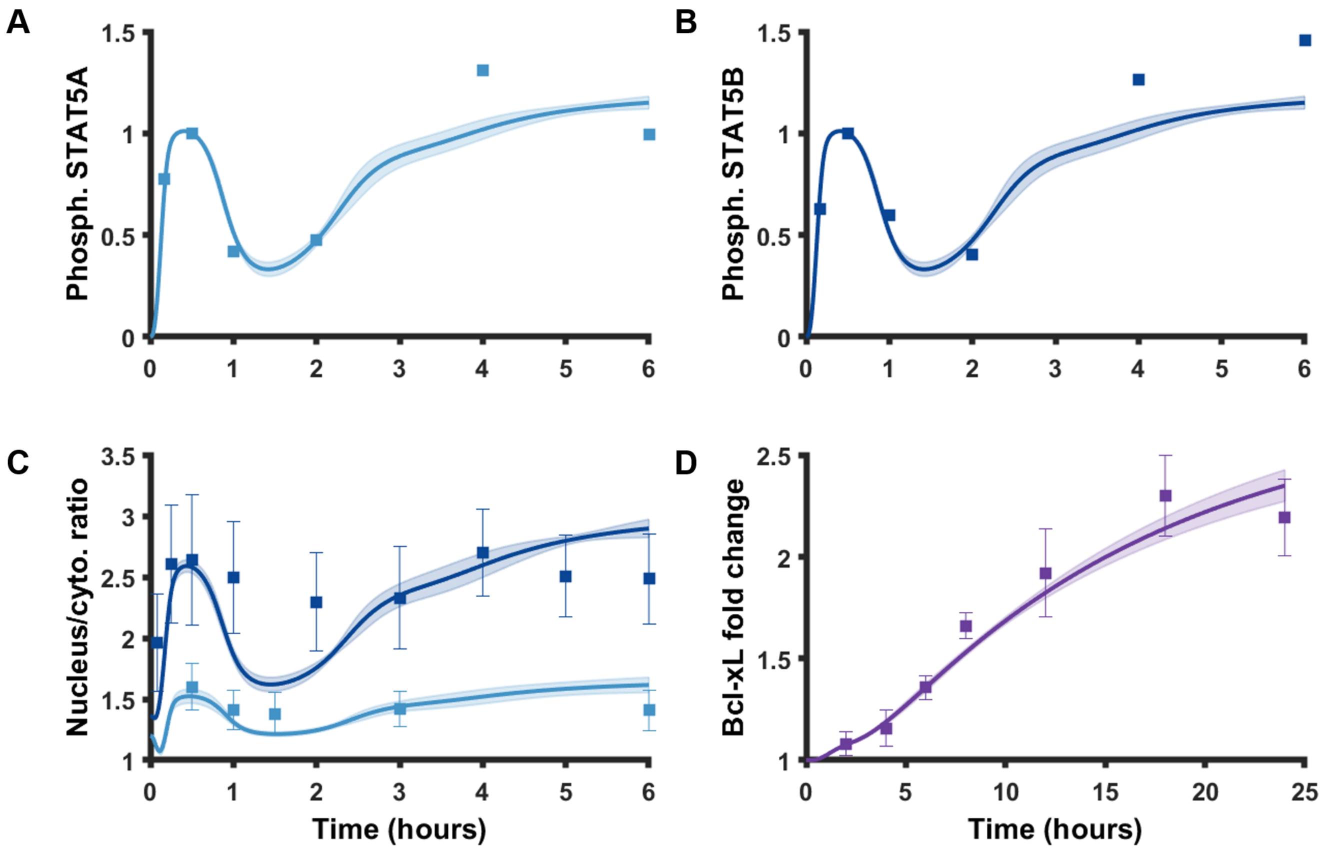

A
B
C
D
Phosph. STAT5A
Phosph. STAT5B
Nucleus/cyto. ratio
Bcl-xL fold change
Time (hours)
Time (hours)

**Figure S7**

a - c

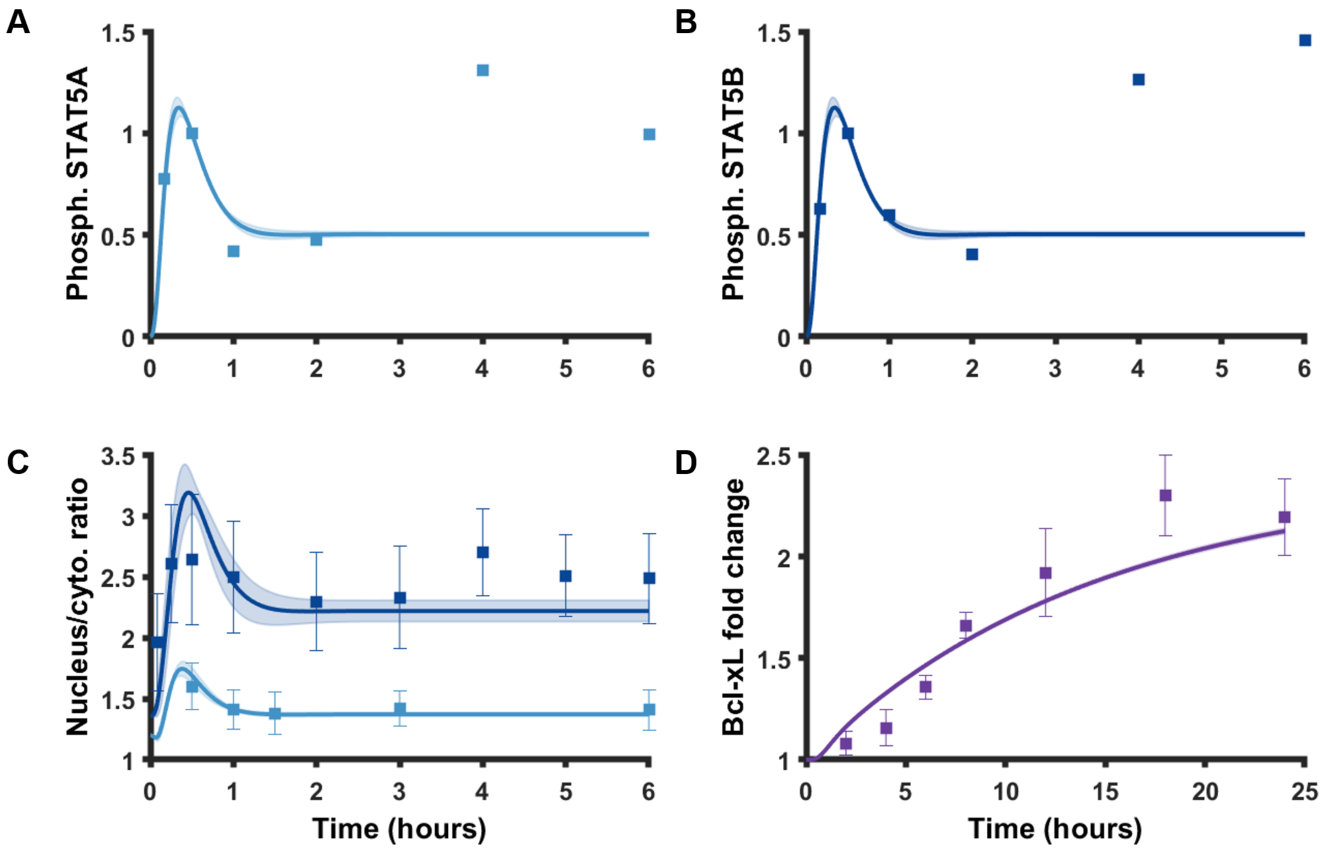
A
Phosph. STAT5A
B
Phosph. STAT5B
C
Nucleus/cyto. ratio
Time (hours)
D
Bcl-xL fold change
Time (hours)

**Figure S8**

- b c

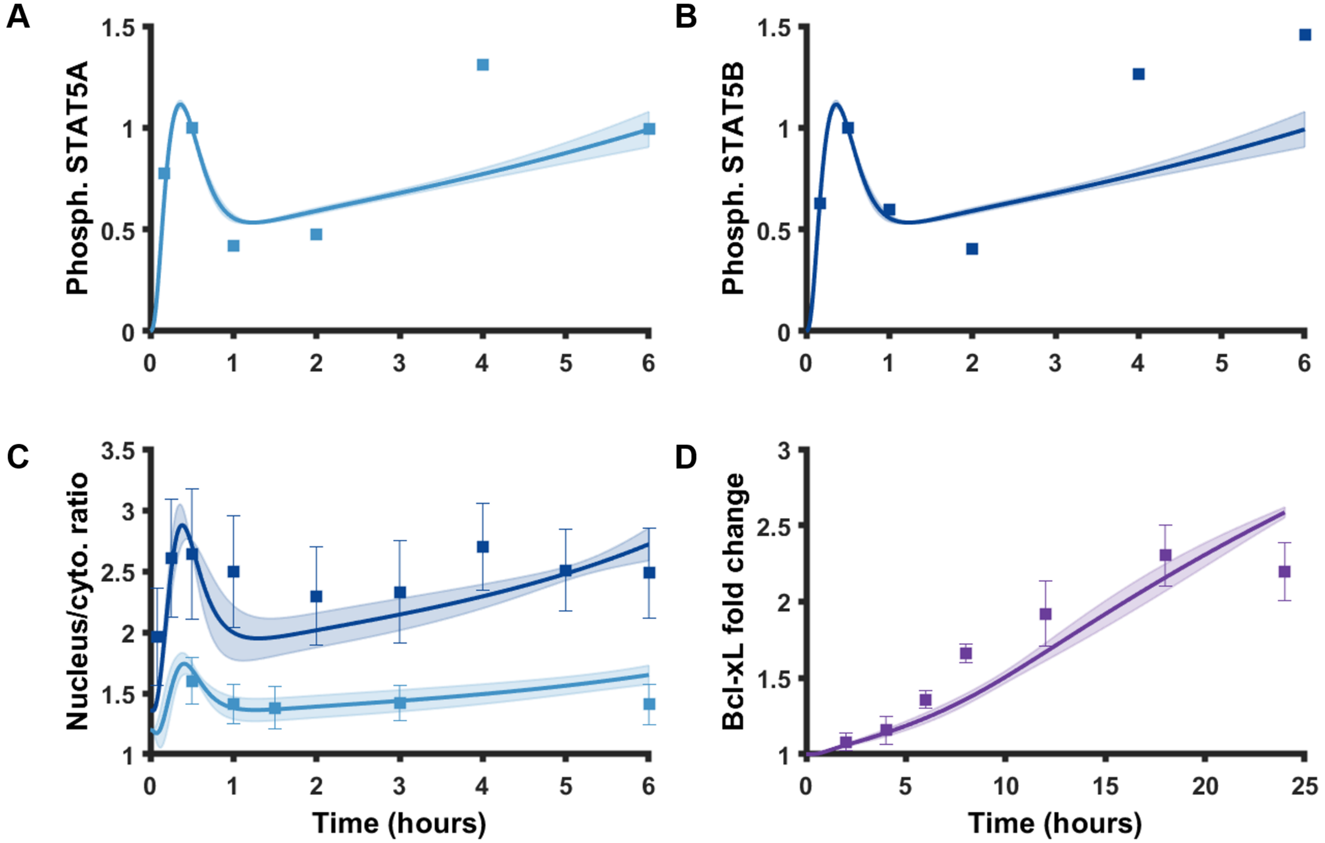
A
B
C
D
Phosph. STAT5A
Phosph. STAT5B
Nucleus/cyto. ratio
Bcl-xL fold change
Time (hours)
Time (hours)

<u>Figure S9: Dose Response Curves for STAT5B</u>

The predicted time course of STAT5B import into the nucleus under various concentrations of PRL ligand, simulated for 60 minutes. The red dotted line emphasizes the values at the 18 minute and 30 minute timepoints, which are plotted in the bar chart in Fig. 7A in order to compare with the experimental data from rat primary beta cells.

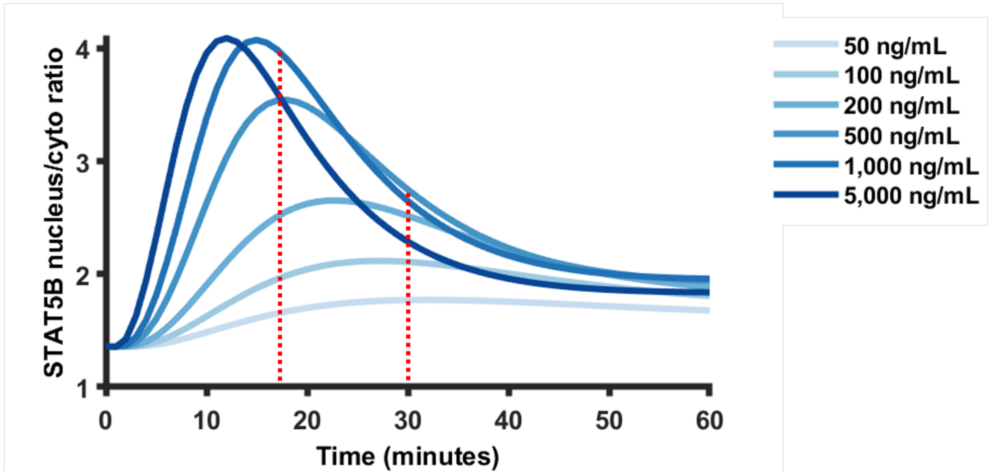

Figure S10: Parameter Value Distributions from Best-Fit Likelihood Estimation

Histograms showing the distribution of each parameter value from the last 8,000 iterations of the fit that had the lowest error during model calibration (Fig. 6). In the first 2,000 iterations, the Bayesian algorithm seeks convergence and in iterations 2,001-10,000 the posterior is sampled for each parameter value.

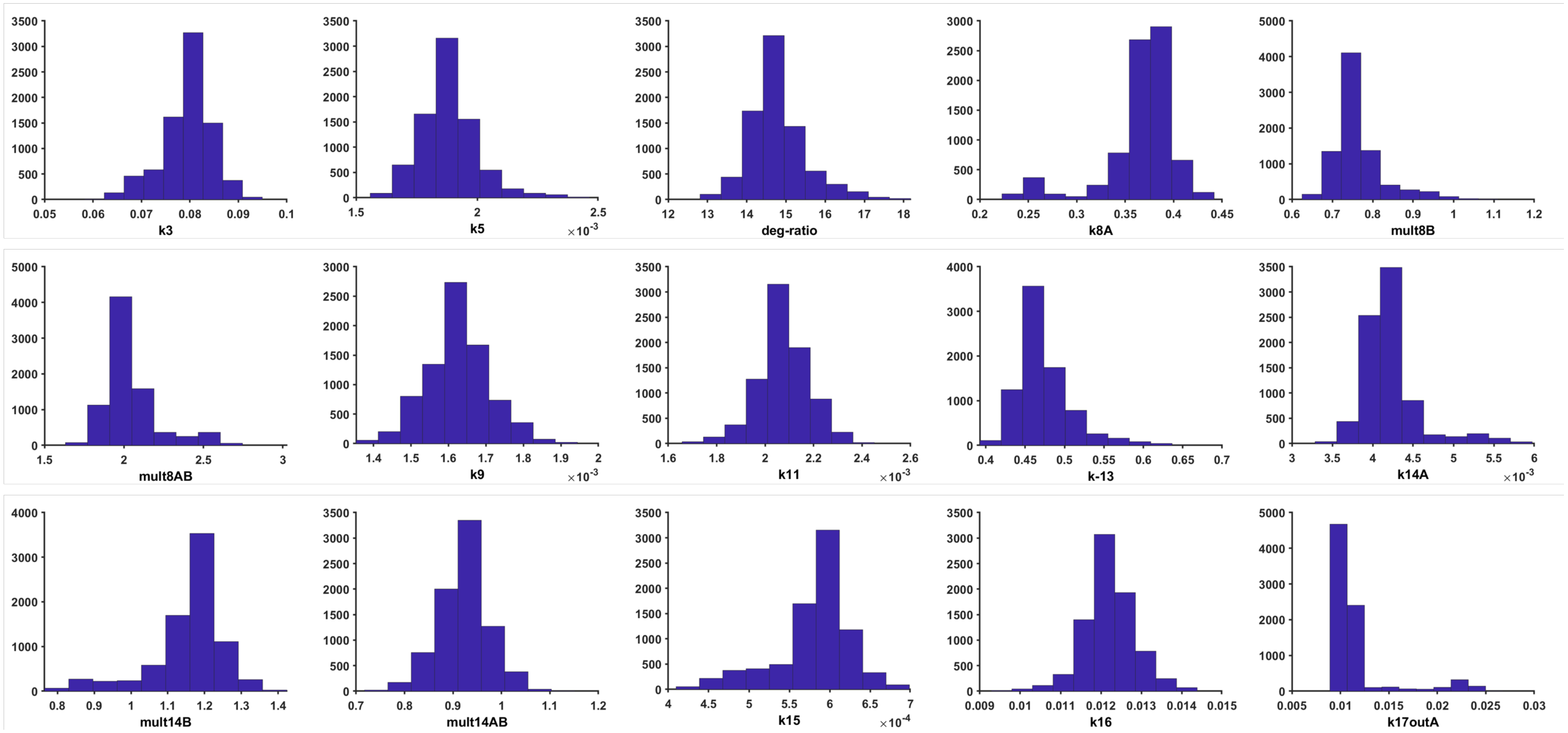

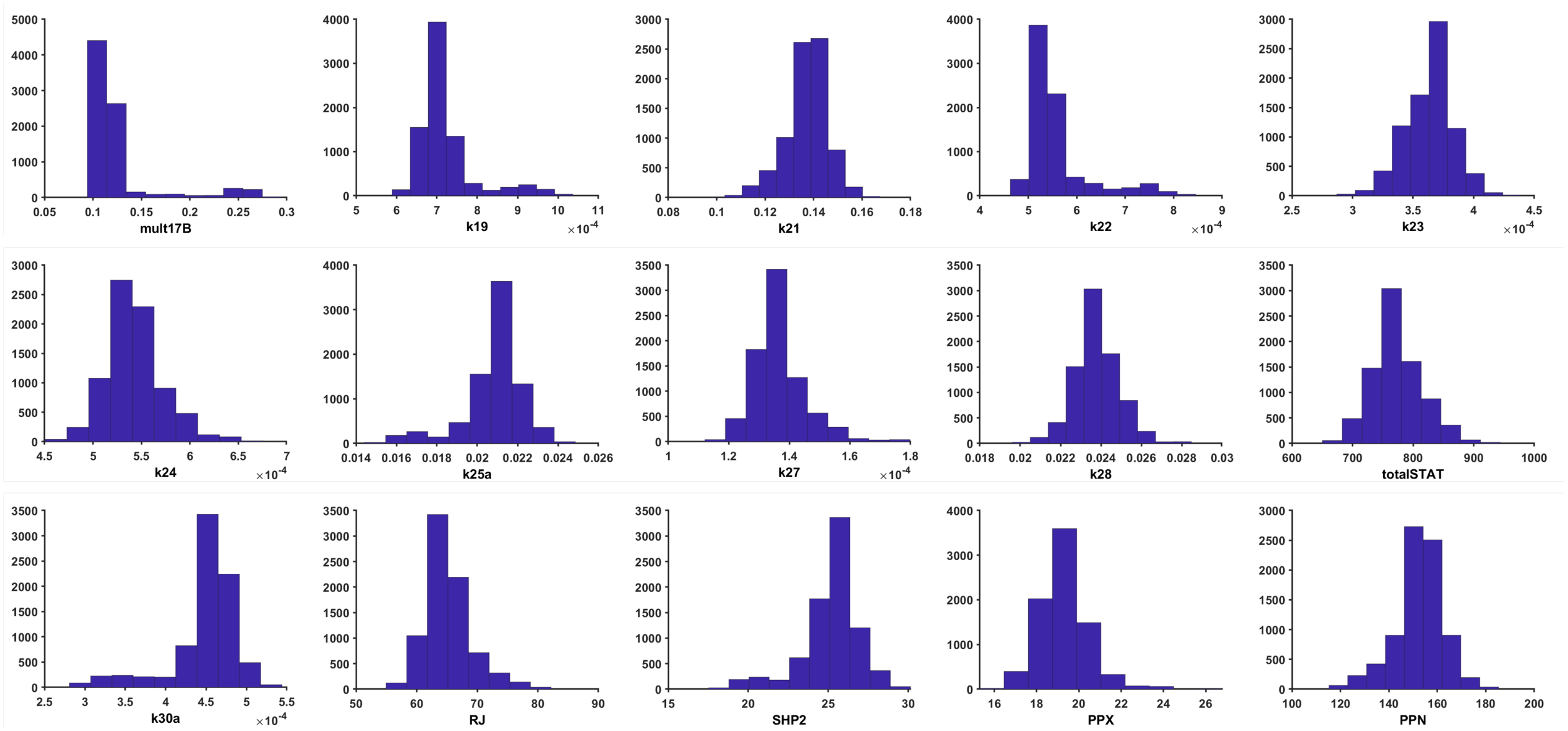
mult17B
k19
×10-4
k21
k22
×10-4
k23
×10-4
k24
×10-4
k25a
k27
×10-4
k28
totalSTAT
k30a
×10-4
RJ
SHP2
PPX
PPN

Figure S11: Parameter Value for Four Best Fits as a Function of Error

Scatter plots for each fitted parameter showing the sum of squared error as a function of the parameter value for the four fits with the lowest error out of the 25 independent fits. In order of lower to higher minimum error: blue, red, yellow, purple. The inter-fit variation in fitted parameter values exceeds the variation within a given fitting. The horizontal spread of the parameter values gives some insight into the importance of the parameter value for the likelihood estimation. For example, for *k5* (second panel), top fit (blue color), the value does not change much as the algorithm converges, indicating the value of *k5* is not affecting the sum of squared errors as much as other parameters

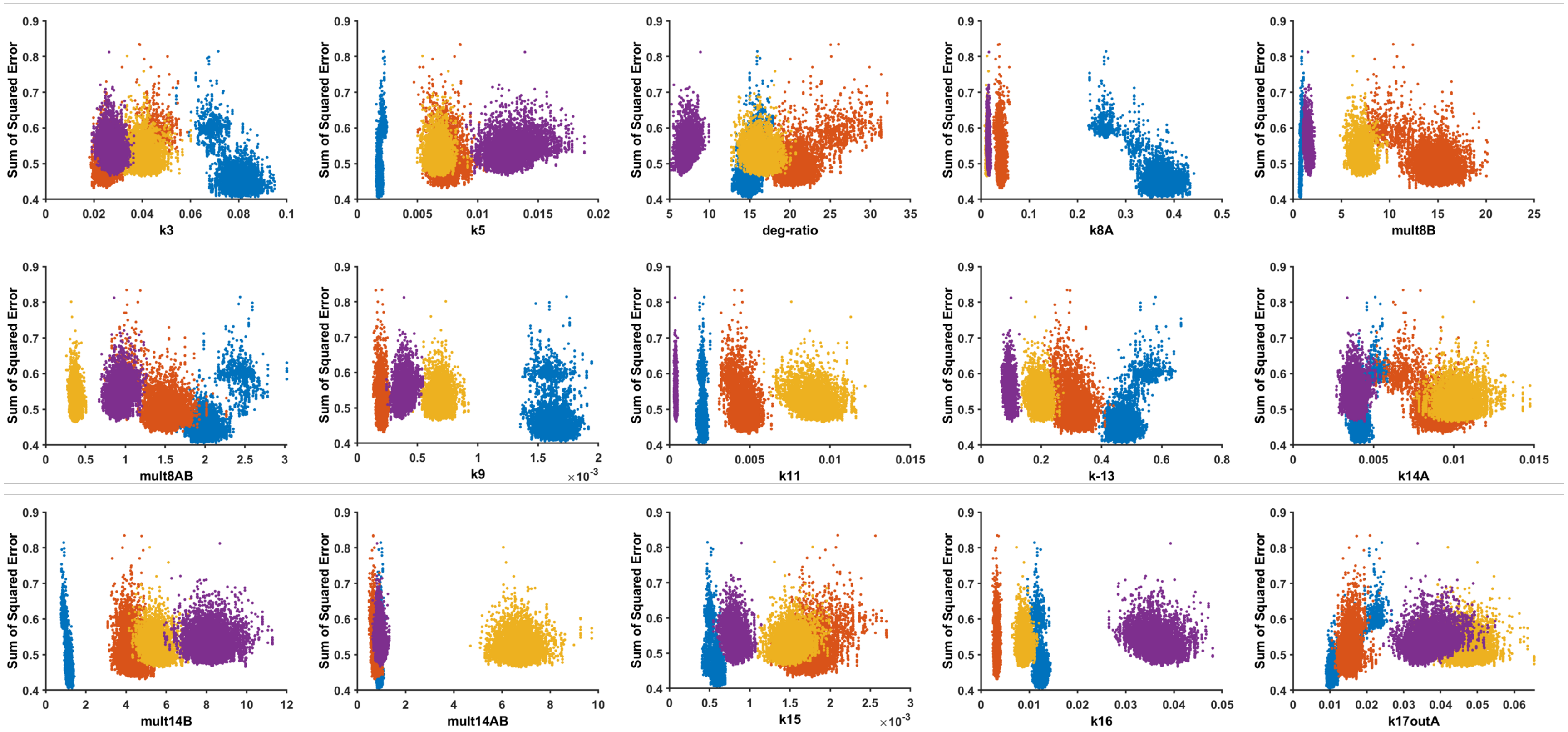

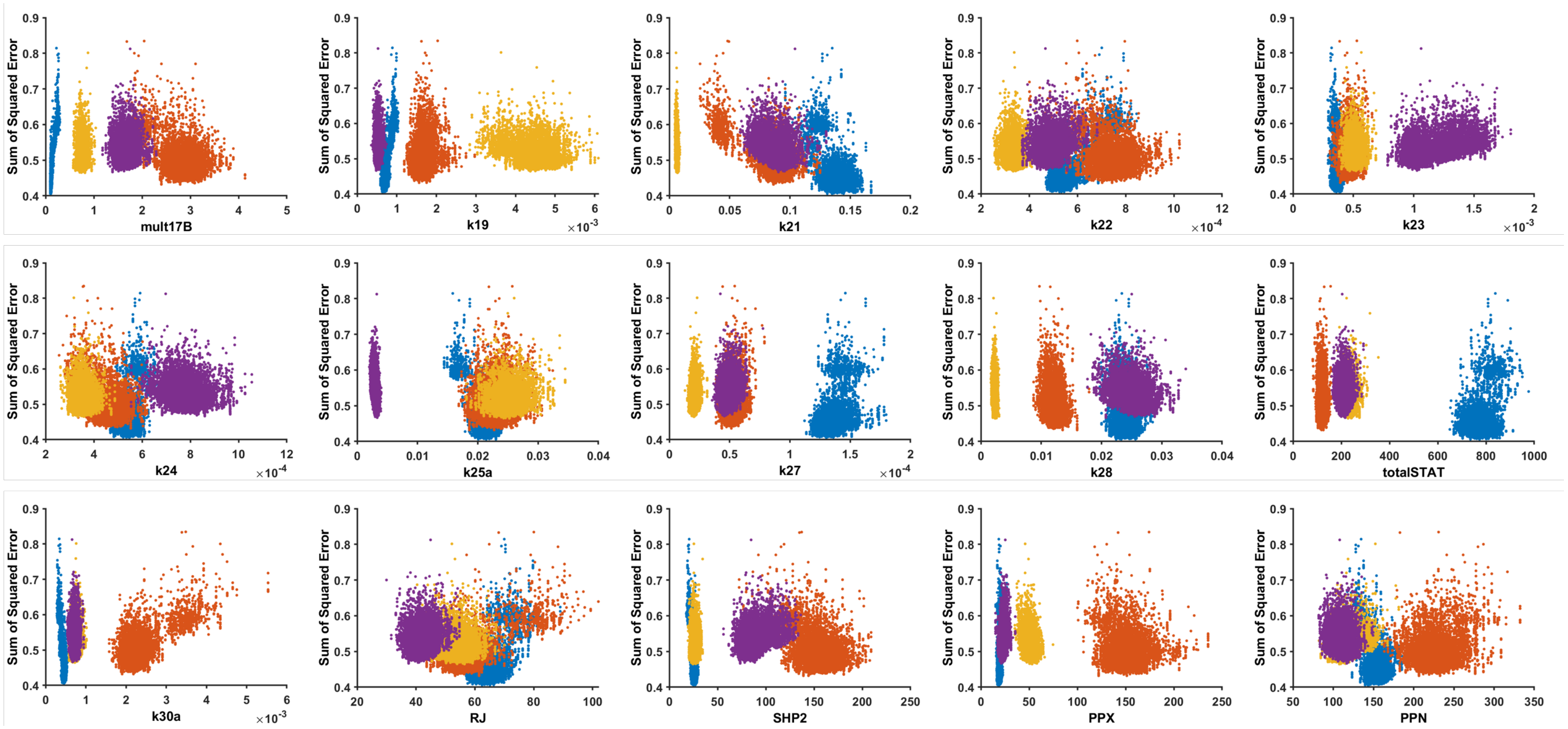

Sum of Squared Error
mult17B
k19
×10^-3
k21
k22
×10^-4
k23
×10^-3
k24
×10^-4
k25a
k27
×10^-4
k28
totalSTAT
k30a
×10^-3
RJ
SHP2
PPX
PPN

Figure S12: Effect of Varying Single Parameters on Model Predictions

Each of the parameters (**Panel A,** *k2*; **Panel B,** *k12*) or initial values (**Panel C,** *PPX*; **Panel D,** *RJ*) were varied one order of magnitude smaller and larger than the fitted value in order to evaluate how perturbations to single parameter values affect model predictions. Lines indicate the mean model prediction from applying the perturbation to all 25 fitted parameter sets. STAT5A predictions are shown separately from STAT5B for ease of viewing. *Dark blue,* STAT5A; *light blue,* STAT5B; *purple,* Bcl-xL. Note the different y-axis limits for Panel D.

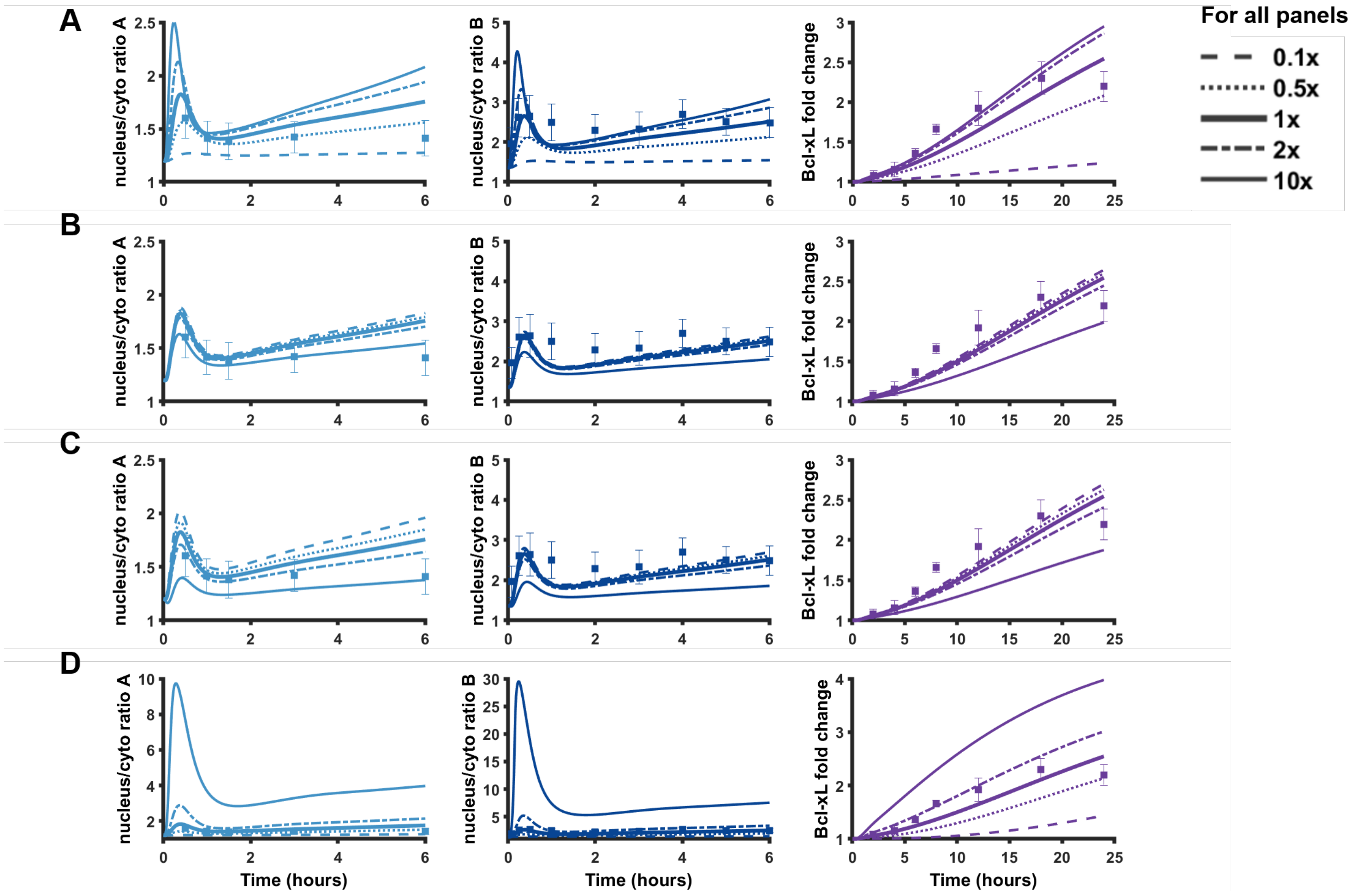
